# Assembly of novel, nuclear dimers of the PI3-Kinase regulatory subunits underpins the pro-proliferative activity of the Cdc42-activated tyrosine kinase, ACK

**DOI:** 10.1101/791277

**Authors:** Natasha S. Clayton, Millie Fox, Jose J. Vicenté-Garcia, Courtney M. Schroeder, Trevor D. Littlewood, Jonathan I. Wilde, Jessica Corry, Kadalmani Krishnan, Qifeng Zhang, Michael J. O. Wakelam, Murray J. B. Brown, Claire Crafter, Helen R. Mott, Darerca Owen

**Affiliations:** Department of Biochemistry, University of Cambridge, 80 Tennis Court Road, Cambridge, CB2 1GA, U.K; GlaxoSmithKline Medicines Research Centre, Gunnels Wood Rd., Stevenage, Herts, SG1 2NY, U.K; The Babraham Institute, Babraham Research Campus, Cambridge, CB22 3AT, U.K; Bioscience, Research and Early Development, Oncology R&D, AstraZeneca, Cambridge, U.K

## Abstract

The tyrosine kinase ACK is an oncogene associated with poor prognosis in human cancers. ACK promotes proliferation, in part, by contributing to the activation of Akt, the major PI3-Kinase effector. We show that ACK also regulates PI3-Kinase directly, via interactions with the PI3-Kinase regulatory subunits. ACK interacts with all five regulatory subunit isoforms and directly phosphorylates p85α, p85β, p55α and p50α on Tyr607 (or equivalent). Phosphorylation of p85β at this residue promotes cell proliferation but, counterintuitively, ACK does not stimulate PI3-Kinase catalytic activity. We show that ACK stabilizes p85α levels by promoting an interaction between the p85 nSH2 domain and pTyr607, protecting p85 from ubiquitination. We demonstrate that ACK interacts with p85α exclusively in nuclear-enriched cell fractions where the increased levels of the regulatory subunits, together with the nSH2-pTyr607 interaction, promote formation of dimeric p85. We postulate that these novel dimers undertake nuclear functions that contribute to Cdc42-ACK driven oncogenesis. We propose that ACK shapes PI3-Kinase signalling by dampening the PIP_3_ response, whilst continuing to drive cell proliferation through Akt activation and hereto unexplored but crucial functions of nuclear dimeric p85. These new regulatory subunit dimers represent a previously undescribed mode of regulation for PI3-Kinase and potentially reveal additional avenues for therapeutic intervention.

## Introduction

The small G protein Cdc42 controls the formation of filopodia by initiating actin remodelling and this process is vital to cell motility. Cdc42 often signals downstream of the master regulator small G protein, Ras, and Cdc42 deregulation results in the progression of several disease states, including tumour metastasis (1). Genetic deletion of *CDC42* in Ras-transformed cells results in reductions in cell proliferation and cell cycle progression, demonstrating the requirement for Cdc42 in Ras-induced transformation (2). Accordingly, overexpression of Cdc42 is seen in several human cancers and is correlated with poor disease outcome (3–5). Although mutations in Cdc42 are rarely observed in human cancers, oncogenic mutations have been characterised in its regulators such as the GEFs Dbl (6) and Asef2 (7) and the GAP DLC1 (8), all of which act to increase the basal level of Cdc42⋅GTP contributing to cellular transformation. Cdc42 participates in both physiological and tumorigenic processes by interacting with effector proteins including the non-receptor tyrosine kinase, activated Cdc42-associated kinase (ACK) (9).

ACK is a ubiquitously expressed non-receptor tyrosine kinase first identified as an effector for Cdc42 (9). The N-terminus of the 120 kDa ACK protein comprises a sterile alpha motif (SAM) domain, nuclear export signal (NES), tyrosine kinase domain, Src homology-3 (SH3) domain and a Cdc42/Rac-interactive binding (CRIB) motif. The proline-rich C-terminus of ACK includes a clathrin binding region, an epidermal growth factor (EGF) receptor-binding domain (EBD) and a ubiquitin association domain (UBA) (Figure 1A).

**Figure 1:**
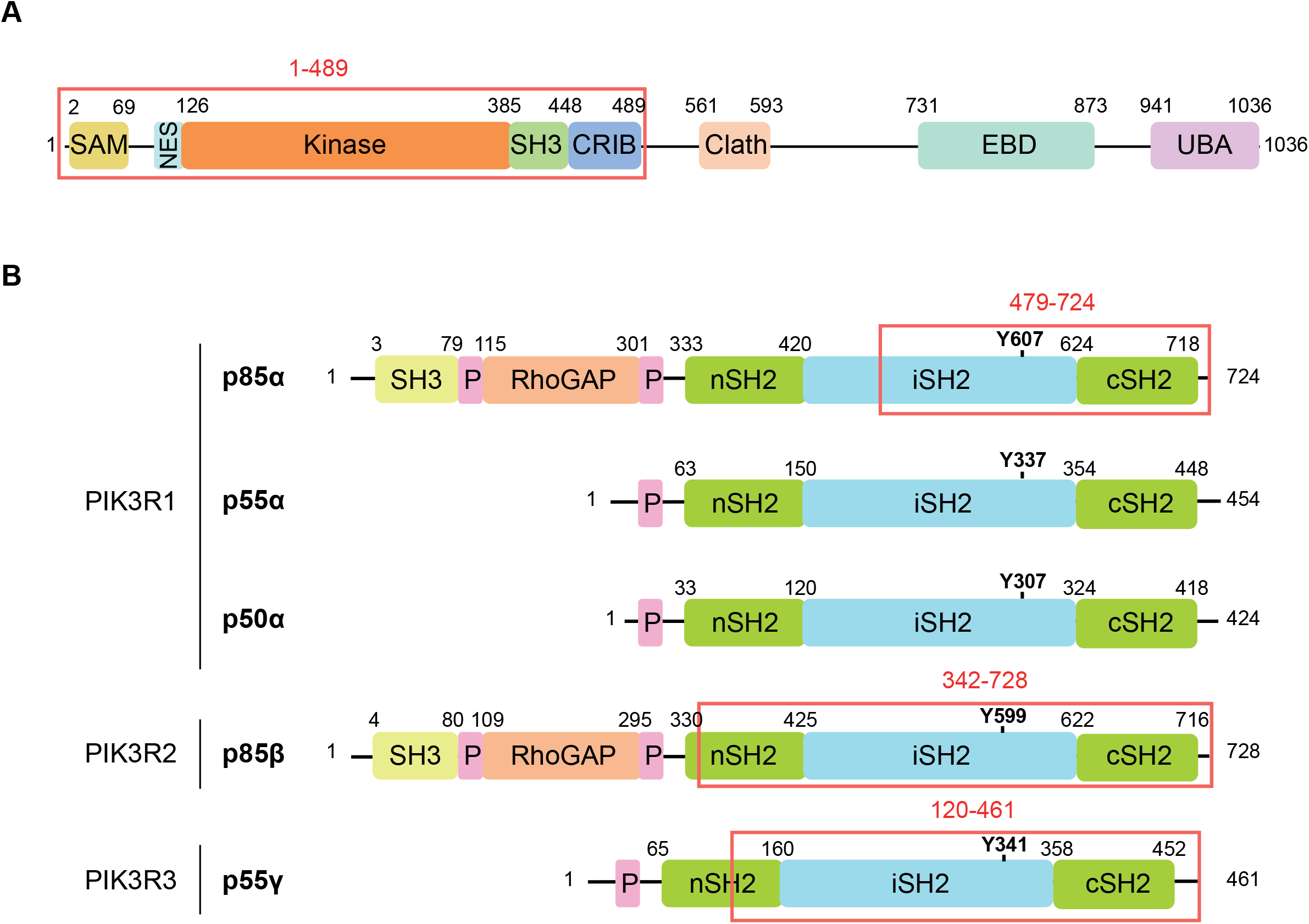
Domain architecture of ACK and the p85 regulatory subunit with a summary of the yeast two hybrid results. (A) Domain architecture of ACK. SAM: sterile alpha motif domain, NES: nuclear export signal, SH3: Src homology-3 domain, CRIB: Cdc42/Rac-interactive binding motif, Clath: clathrin binding region, EBD: EGFR binding domain, UBA: ubiquitin association region. The region in the N-terminus of ACK used for the yeast-2-hybrid screen is indicated by the red box. (B) Domain architecture of class Ia PI3-Kinase regulatory subunits. SH3: Src homology-3 domain, P: proline-rich sequences, RhoGAP: region with sequence homology to Rho family GTPase-activating domains, SH2: Src homology-2 domain, nSH2: N-terminal SH2 domain, cSH2: C-terminal SH2 domain, iSH2: inter-SH2 region. Regions involved in interactions with ACK, as identified by the yeast-2-hybrid screen, are indicated by red boxes. The ACK phosphorylation site in p85α and the equivalent tyrosine residues in p50α, p55α, p85β and p55γ are shown in bold.

In the past decade, ACK has emerged as a prospective therapeutic target in multiple human cancers. A genomic study, which sought to identify oncogenic ‘driver’ mutations within the protein kinase gene family in 210 diverse cancers, identified four such mutations in ACK (10), ranking ACK within the top ~5% of kinases driving cancer. Amplification or mutation of the gene encoding ACK is common in aggressive lung, ovarian and hormone refractory prostate tumours, and increased levels of ACK mRNA correlate with poor patient prognosis (10, 11). Expression of activated ACK in LNCaP cells increases the ability of the cells to form tumours in nude mice (12). Ultimately, the successful development of therapeutic inhibitors that down-regulate ACK signalling will rely on a detailed understanding of the pathways that feed into ACK activation and the pathways that are activated downstream of ACK.

Although ACK is implicated in tumourigenesis, relatively little is known about its cellular substrates or interacting proteins, leaving the full molecular details of its function unknown. Headway has been made in understanding the role of ACK in prostate cancer. ACK phosphorylates the tumour suppressor Wwox, leading to its ubiquitination and degradation in the later stages of prostate cancer progression (12). ACK also phosphorylates histone H4, which ultimately leads to up-regulation of the Androgen Receptor (AR) (13) and, in a co-ordinated manner, ACK phosphorylates the AR directly, activating its transcriptional activity (14). Both activities contribute to progression to the castration-resistant phase of prostate cancer.

We have identified the class Ia PI3-Kinase regulatory subunit isoforms as novel ACK interacting proteins. PI3-Kinases catalyse the formation of the second messenger PIP_3_ from PIP_2_, which leads to the activation of pathways that promote cell proliferation and survival. Class Ia PI3-Kinases consist of a catalytic subunit (p110), of which there are three isoforms (p110α, β and δ), bound to a regulatory subunit (p85), which exists in five isoforms: p85α, p85β, p55α, p50α and p55γ. The five regulatory isoforms have a similar domain organisation at the C-terminus (Figure 1B), with a proline rich sequence and two Src homology-2 (SH2) domains, separated by a coiled-coiled domain termed the inter-SH2 (iSH2) region. The larger regulatory subunit isoforms, p85α and p85β, also possess additional functional domains at the N-terminus, including a Src homology-3 (SH3) domain, proline-rich sequences and a RhoGAP domain (often referred to as the BH domain), which shows sequence homology to the GTPase-activating protein (GAP) domain of the breakpoint cluster region protein, Bcr. p110α is the most commonly mutated kinase in the human genome (15) and cancer-associated mutations or alterations in gene copy number have been identified for the genes encoding each of the three catalytic subunit isoforms (*PIK3CA*, *PIK3CB* and *PIK3CD)* and the genes encoding each of the five regulatory subunit isoforms (*PIK3R1*, *PIK3R2* and *PIK3R3*) (16). In fact *PI3KR1* is the 11^th^ most mutated gene across a subset of 20 human cancers (17). Over-activation of PI3-Kinase signalling in cancer can also result from mutations that reduce the activity of the PTEN phosphatase, which counteracts PI3-Kinase activity by dephosphorylating PIP_3_ back to PIP_2_, or from mutations that activate the major downstream effector kinase, Akt.

ACK also modulates PI3-kinase signalling pathways by directly phosphorylating Akt, which promotes Akt membrane localization and subsequent phosphorylation, resulting in its activation (18). Here, we report that ACK interacts with other key components of the PI3-Kinase pathway, the PI3-Kinase regulatory subunits. ACK interacts with all five regulatory subunits of PI3K, phosphorylating four of them in the iSH2 region. This pTyr serves to stabilise p85 by preventing ubiquitination and proteasome degradation, leading to increased cellular levels of the regulatory subunits. The mechanism underpinning this stabilization involves an interaction between the pTyr and the nSH2 domain of the regulatory subunit. This interaction *in trans* supports formation of a novel dimeric form of p85, which is favoured due to increased levels of the regulatory isoforms, leading to a pool of p110-free p85. This new mechanism of dimerization mediated by sites in the C-terminal regions of the p85 isoforms is also possible in the shorter isoforms, permitting multiple homo and hetero dimer combinations. We further demonstrate that the interaction between ACK and p85 occurs exclusively in the nucleus, where the resulting phosphodimers are also mainly located.

We show that the proliferative advantage conferred by ACK is driven, at least in part, by its effects on the regulatory subunits and propose that ACK modulates the PI3-Kinase response at multiple nodes including hereto unexplored but crucial functions of novel C-terminally mediated p85 dimers in the nucleus. These new regulatory subunit dimers represent a previously unidentified mode of regulation for PI3-Kinase and potentially reveal additional avenues for therapeutic intervention.

## Materials and Methods

### Yeast-2-hybrid screen

ACK (1-489) was cloned into pENTR/D-TOPO and then transferred into destination vectors GpYTH9 and GpYTH16, using Gateway recombination. This produced recombinant yeast expression bait constructs with the ACK coding region in frame with the DNA binding domain of Gal4. The GpYTH16 construct was used to assess trans-activation in *S. cerevisiae* using a β-galactosidase assay, while the GpYTH9 construct was linearized with *Xba*1 and integrated into the yeast strain Y190::ADE2 (MATa, ura3-52, his3-Δ200, lys2-801, trp1-901, leu2-3, 112, gal4Δ, gal80Δ, cyh^r^2, LYS2::GAL1_UAS_-HIS3_TATA_-HIS3, MEL1 URA3::GAL _UAS_-GAL1_TATA_-lacZ, GAL2_UAS_-GAL2 _TATA_-ADE2) for screening. Transformants were selected on media lacking tryptophan. The resulting strain was transformed with a Matchmaker Human Foetal Brain cDNA library (Clontech), where prey cDNAs are cloned in frame with the activation domain of Gal4 in pACT2. Transformants were selected on media lacking leucine but containing X-gal (5-Bromo-4-Chloro-3-Indolyl β-D-Galactopyranoside). Large blue colonies were picked onto fresh selective plates and then reassayed for β-galactosidase activity. Yeast colonies were transferred onto Whatman No.54 filter papers and lysed by freeze thawing. Each filter was then immersed in 60 mM NaH_2_PO_4_, 40 mM NaH_2_PO_4_, 10 mM KCl, 0.1 mM MgSO_4_, 40 mM β-mercaptoethanol, 1 mg/mL X-gal, pH 7.0 and incubated at 37 ºC for 4 h. Filters were air-dried and checked for blue colouration. Library plasmids isolated from the β-galactosidase-expressing Leu^+^ yeast, were recovered and their sequences analysed.

### Plasmids and site-directed mutagenesis

pXJ-HA-ACK was a kind gift from Prof Ed Manser (ICMB, Singapore). Full-length human p85α, p50α, p55α and p55γ and mouse p85β cDNA (IMAGE) were amplified by PCR, cloned into pENTR/D-TOPO and then transferred into the mammalian expression vector pDEST12.2-FLAG and the bacterial expression vectors pDEST15 and pETG-41A using Gateway technology. p85β cDNA was also cloned into the lentiviral vector, pSLIK-hygro. The coding sequences of wt and caACK were digested from pXJ-HA constructs *Eco*RI/*Xho*I and ligated into the retroviral expression vector, pBABE-puro. Site-directed mutagenesis was performed using a QuikChange Lightning Multi Site-Directed Mutagenesis Kit (Agilent Technologies).

### Antibodies

The following antibodies were used for immunoblotting: anti-FLAG-HRP (A8592, Sigma Aldrich), anti-V5 (46-0705, Invitrogen) and anti-GST (27-4577-01, GE Healthcare); anti-p-Tyr-HRP (sc-7020), anti-p85 (sc-423), anti-Hsp56 (sc-1803), anti-GAPDH (sc-47724) and anti-HA-HRP (sc-7392) were purchased from Santa Cruz Biotechnology; anti-ACK (161-175 and 07-757) and anti-ACK p-Tyr284 (09-142) were purchased from Millipore; anti-Histone H3 (ab1791) and anti-pY607 p85 (ab182651) were purchased from Abcam. The HRP-conjugated secondary antibodies donkey anti-mouse (sc-2318) and donkey anti-rabbit (sc-2317) were purchased from Santa Cruz Biotechnology. Anti-FLAG (F3165, Sigma Aldrich) was used for immunoprecipitation.

### Cell Lines

HEK293T cells were cultured in DMEM supplemented with 10% FBS and transfected using Lipofectamine 2000. To generate HEK293T cell lines stably expressing HA-wtACK or HA-caACK, Phoenix-AMPHO cells were transfected with pBABE-puro-wtACK or caACK. Retroviral supernatant was collected after 72 h and added to HEK293T cells for 16 h with 4 ng/µL polybrene. Infected cells were selected using 1 µg/mL puromycin dihydrochloride. To generate HEK293T cell lines stably expressing FLAG-wt p85β/p85β Y599F, HEK293T cells were transfected with 8 µg pSLIK-hygro-FLAG-wt p85β/p85β Y599F and 4 µg each of the packaging vectors pCMV-VSV-G and pCMV-∆R8.2 dvpr. After 16 h, viral supernatant was added to recipient HEK293T cells for 16 h with 4 ng/µL polybrene. Infected cells were selected using 150 µg/mL Hygromycin B. Transgene expression was confirmed by immunoblotting. Control cell lines were generated in the same way using empty vector but were expanded as a polyclonal cell line.

### Cell Proliferation Assays

Cells were seeded into 6-well plates at a density of 3×10^4^ cells per well and incubated at 37 °C, 5% CO_2_. Assays using ACK stable cell lines were performed in DMEM + 0.5% FBS. Assays using p85β cell lines were performed in DMEM + 10% FBS. p85β expression was induced using 1 mg/ mL doxycycline. Cells were trypsinized and dispersed using a syringe before being counted using a Countess automated cell counter at specific timepoints.

### Co-immunoprecipitation

Small-scale co-immunoprecipitations were performed using 1 µg antibody cross-linked to Protein G Dynabeads (Life Technologies) with dimethyl pimelimidate (DMP). HEK293T cells were grown in 10 cm dishes to 70% confluence and then transfected for 40 h. Cells were then washed in cold PBS and lysed on ice in 500 µL mammalian cell lysis buffer (50 mM Tris-HCl pH 7.5, 150 mM NaCl, 1 mM EDTA, 1 mM sodium orthovanadate, 1 mM β-glycerophosphate disodium salt hydrate, 1X Mammalian Protease Inhibitor Cocktail (Sigma Aldrich, P8340), 1% Triton X-100). Lysates were centrifuged at 17000 g for 20 min at 4 °C and supernatants were pre-cleared by incubation with 50 µL Dynabeads at 4 °C for 1 h. Once cell lysates had pre-cleared, 500 µL of each lysate was rotated with 50 µL antibody-bound beads for 1h. The beads were then washed three times with mammalian cell lysis buffer and protein complexes were eluted using 20 µL 2X LDS sample buffer (Life Technologies) mixed with 20 µL PBS.

### Immunoblotting

Proteins were transferred to Immobilon-P PVDF membrane (Millipore) using an X-Cell II™ Blot Module (Life Technologies) following manufacturer’s instructions. Membranes were blocked in 10% milk dissolved in PBS-0.1% Tween. Protein bands were visualized using enhanced chemiluminescence.

### Cell fractionation

Cells were grown in 10 cm dishes to 70% confluence and then transfected for 40 h. Cells were then washed in PBS, resuspended and pelleted by centrifugation at 500 g for 20 min at 4 °C. Cell pellets were then resuspended in 500 µL cold hypotonic buffer (20 mM Tris-HCl pH 7.4, 10 mM NaCl, 3 mM MgCl_2_) and incubated on ice for 15 min. 25 µL NP-40 Alternative was then added and cells were lysed by vortexing for 10 sec. Samples were then centrifuged at 500 g for 20 min at 4 °C to separate the cytoplasmic fraction from intact nuclei and other subcellular structures. The supernatant (cytoplasmic fraction) was removed and stored on ice. The pellet was washed with 500 µL cold hypotonic buffer and then resuspended in 500 µL cell extraction buffer (100 mM Tris-HCl pH 7.4, 100 mM NaCl, 1 mM EDTA, 1 mM EGTA, 0.1% SDS, 1 mM NaF, 2 mM Na_3_VO_4_, 1% Triton X-100, 10% glycerol, 0.5% deoxycholate, 20 mM Na_4_P_2_O_7_, 1 mM PMSF, 1X Mammalian Protease Inhibitor Cocktail. Pellets were incubated on ice for 30 min with vortexing every 10 min. Insoluble debris was cleared from the nuclear-enriched fractions by sonication and centrifugation at 17000 g for 30 min at 4 °C. Cytoplasmic and nuclear-enriched fractions were then either analysed by SDS-PAGE gel directly or used in co-immunoprecipitation studies.

### Quantification of PIP_3_

HEK293T cells were seeded onto 6-well plates at a density of 1×10^6^ cells per well and incubated at 37 °C, 5% CO_2_ for 24 h before transfection with the appropriate construct for 24 h. Cells were then serum starved for 16 h before stimulation with 10 nM EGF for 5 min. Cells were washed with ice cold PBS, harvested and frozen in liquid nitrogen. Phosphoinositides were extracted as previously described and samples were quantified for total cellular PIP_3_ and PIP_2_ by the mass spectrometry (19).

### Expression and purification of recombinant regulatory subunits and domains

Full-length human p85α, p50α, p55α, mouse p85β, p85α nSH2 (325-430), p85α cSH2 (614-720) and p85β cSH2 (603-712) were expressed in *E. coli* as GST fusion proteins. Expression was induced by 1 mM IPTG for 16 h at 20 °C. Cells were harvested and resuspended in MTPBS (150 mM NaCl, 16 mM Na_2_HPO_4_, 4mM NaH_2_PO_4_ pH 7.3) supplemented with SIGMAFAST Protease Inhibitor Cocktail Tablet (EDTA-free) and 1 mM PMSF, and lysed using an Emulsiflex at 10000 p.s.i. Lysates were centrifuged at 18000 g for 40 min at 4 °C and the supernatant applied to glutathione-agarose beads for 2 h at 4 °C. After washing, GST-full-length p85 fusion proteins were eluted from the beads with 10 mM reduced glutathione. GST-SH2 fusion proteins were subject to HRV 3C cleavage whilst immobilized on GST beads in MTPBS, 1mM DTT, 1mM EDTA. Following cleavage, the SH2 domains were further purified by size exclusion chromatography on an S75 column (GE Healthcare).

Full-length human p55γ and p85β nSH2 (312-425) were expressed as fusion proteins with N-terminal His_6_ and MBP tags. Expression of p55γ fusion protein was induced with 1 mM IPTG for 3 h at 30 °C. Expression of p85β nSH2 fusion protein was induced with 1mM at 37 °C for 5 h. Cell pellets were resuspended in lysis buffer (20 mM Tris-HCl pH 7.9, 500 mM NaCl) supplemented with Protease Inhibitor Cocktail (Sigma Aldrich, S8830) and 1 mM PMSF. Cells were lysed as described above.

Cleared lysate was loaded onto a Ni-NTA column and proteins eluted using and imidazole gradient. His_6_-MBP-p85β nSH2 fusion was subject to HRV 3C cleavage and further purification by size exclusion chromatography as described previously.

### Expression and purification of His6-ACK (110-489) and GST-ACK (1-489)

ACK residues 110-489 were cloned into pFastBac-HT and recombinant bacmid was generated, using *E. coli* DH10Bac, to allow expression of His_6_-ACK (110-489) using the baculovirus/Sf9 cell system. ACK residues 1-489 were cloned into pDEST15 using Gateway technology. GST-ACK (1-489) was expressed in BL21 Rosetta 2(DE3) cells. His_6_-ACK (110-489) and GST-ACK (1-489) were purified as previously described (20).

### *In vitro* kinase assays

His_6_-ACK (110-489) or GST-ACK (1-489) was incubated for 10 min at 30 °C in kinase buffer (20 mM Tris-HCL pH 7.5, 10 mM MgCl_2_, 0.5 mM DTT, 0.1 mM sodium orthovanadate) supplemented 0.5 mM ATP pH 7.5 to promote autophosphorylation. For *in vitro* kinase assays, 0.3 mM autophosphorylated His_6_-ACK (110-489) or GST-ACK (1-489) was added to kinase buffer with 30 mM substrate in a final reaction volume of 50 μL and incubated for 30 min at 30 °C.

### Phosphorylation site mapping using LC-MS/MS

Samples were resolved by SDS-PAGE and visualized using InstantBlue (Expedeon). Bands of interest were excised from the gel, destained, reduced using DTT and alkylated using iodoacetamide. The protein was then digested with trypsin overnight at 37 °C and then loaded onto a nanoAcquity UPLC (Waters Corp., Milford, MA) system and an LTQ Orbitrap Velos hybrid ion trap mass spectrometer (Thermo Scientific, Waltham, MA) for analysis. The peptides were first separated by reverse-phase liquid chromatography (LC) using a Waters reverse-phase nano column (BEH C18, 75 mm i.d. x 250 mm, 1.7 mm particle size) at flow rate of 300 nL/min after pre-loading onto a pre-column (Waters UPLC Trap Symmetry C18, 180 mm i.d. x 20 mm, 5 mm particle size) in 0.1% formic acid for 5 min at a flow rate of 5 mL/min. The peptides were then eluted from the pre-column and loaded onto the nanoAcquity analytical column. The LC eluant was sprayed into the mass spectrometer using a New Objective nanospray source. The m/z values of eluting ions were measured in the Orbitrap Velos mass analyzer, set at a resolution of 30000. Data dependent scans (top 20) were then employed to automatically isolate and generate fragment ions by collision-induced dissociation in the linear ion trap, resulting in the generation of MS/MS spectra. Ions with charge states of 2+ and above were selected for fragmentation. The MS/MS data were processed using Protein Discoverer (version 1.3, ThermoFisher). Briefly, all MS/MS data files were submitted to the Mascot search algorithm (Matrix Science, London UK) and searched against the PI3-Kinase regulatory subunit 1 sequence, using a fixed modification of carbamidomethyl and variable modifications of oxidation (M) and phospho (Ser,Thr,Tyr). Sites of phosphorylation were then verified manually. The peptide and fragment mass tolerances were set to 25ppm and 0.8 Da, respectively. A significance threshold value of p<0.05 and a peptide cut-off score of 20 were also applied.

### Fluorescent Polarization Assays

p85α and CD28 peptides were purchased from Biomatik; p85β peptides were purchased from Cambridge Research Biochemicals. Peptide sequences were as follows: peptides based on p85α, [5-FAM]-TEDQ-[pY]-SLVEDE and [5-FAM]-TEDQYSLVEDE; peptides based on p85β, [5-FAM]-TEDQ-[pY]-SLMEDE and [5-FAM]-TEDQYSLMEDE; CD28 peptide, [5-FAM]-SD-[pY]-MNMTP. Fluorescence polarization experiments were measured on a BMG Labtech Pherastar fluorimeter at 298 K with excitation 485 nm and emission 520 nm. Solutions were made up in black flat-bottom 384 well plates (Corning) to a final volume of 30 µL. A peptide titration was used to determine the appropriate concentration of peptide for binding assays. CD28, p85α and phospho-p85α were used at 20nM. p85β and phospho-p85β were used at a final concentration of 40nM. Anisotropy was calculated from polarization measurements in MARS analysis software and data fitted to a single site binding isotherm in Prism.

## Results

### Identification of the interaction between ACK and the PI3-Kinase regulatory subunits

A yeast-2-hybrid screen was performed to identify binding partners for ACK. A yeast strain, containing integrated GpYTH16-ACK(1-489) as bait, was screened against a pACT2-based human adult brain cDNA library, expressing prey proteins fused to the GAL4 activation domain. A human brain library was chosen as ACK was initially isolated from a human hippocampal expression library (9) and, although ACK is ubiquitously expressed, it shows high expression levels in the brain (21). 169 β-galactosidase positive colonies were obtained, of which 114 remained after re-screening. Sequence analysis of the positive colonies revealed three isoforms of the PI3-Kinase regulatory subunit, p85α, p85β and p55γ, as the top hits for putative, novel ACK interacting proteins.

The yeast-2-hybrid screen detected an interaction between the N-terminal half of ACK (1-489) and p85α (479-724), p85β (342-728) and p55γ (120-461) (Figure 1A and B), all of which included the C-terminal SH2 domain and most of the inter-SH2 region. The p85α splice variants, p50α and p55α, were not identified in the screen but as they both contain the C-terminal SH2 domain and inter-SH2 region, we reasoned that ACK would likely interact with these isoforms as well. Co-expression of full-length ACK with all five full-length p85 isoforms individually in HEK293T cells, followed by immunoprecipitation, revealed that ACK could interact with all regulatory subunit isoforms (Figure 2A-E). Isoforms p85α, p85β and p55γ were also found to interact with a kinase-dead ACK mutant, dACK (K158R) (Figure 2A, D and E), suggesting that catalytic activity was not required for these novel interactions.

**Figure 2:**
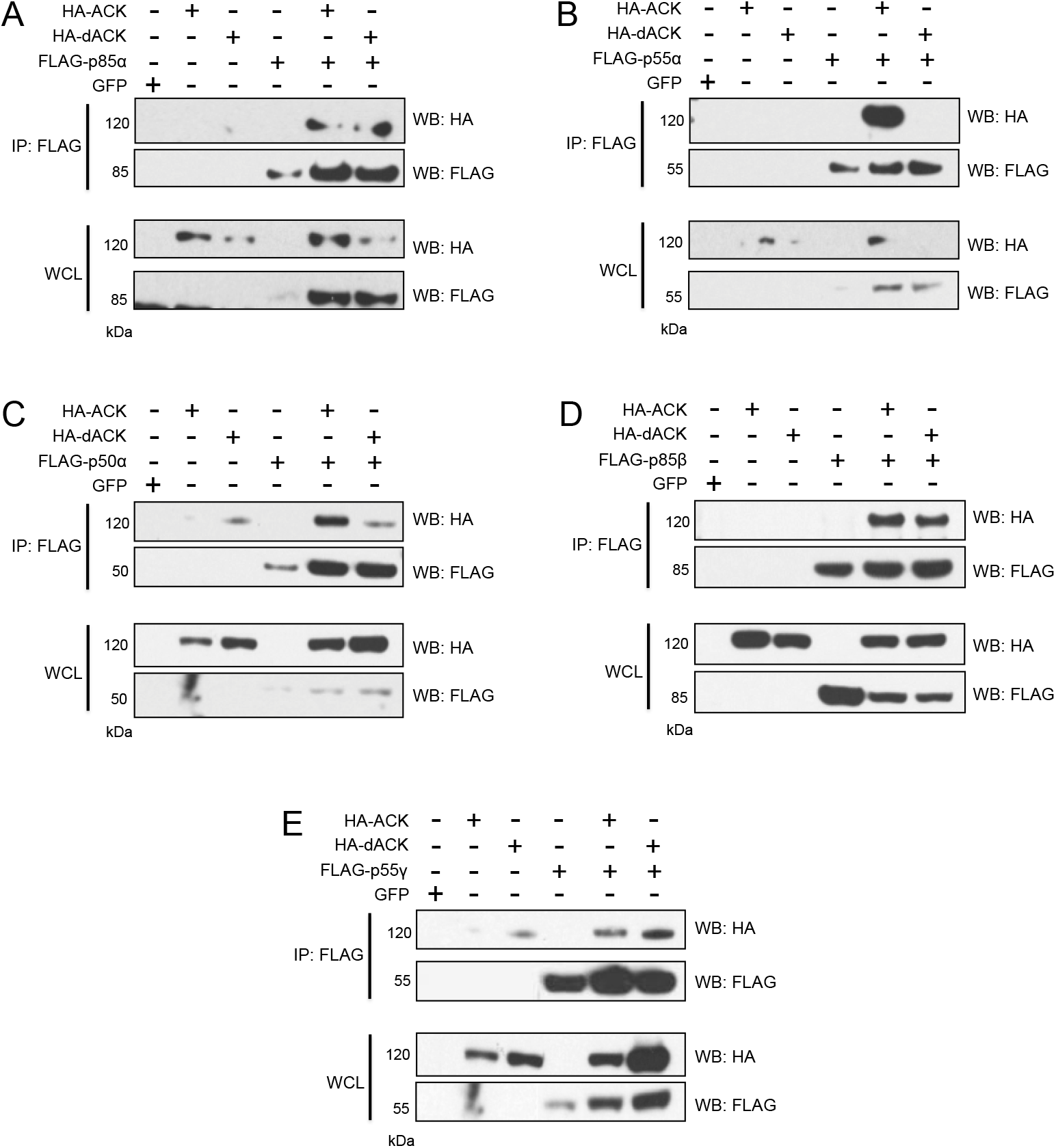
Interaction of ACK with p85α, p55α, p50α, p85β and p55γ in HEK293T cells. HA-ACK and HA-dACK (a kinase dead mutant, K158R) were expressed alone and with the FLAG tagged PI3-Kinase regulatory subunit isoforms p85α (A), p55α (B), p50α (C), p85β (D) and p55γ (E). Expression of the recombinant proteins in whole cell lysate (WCL) is shown in the bottom panels. PI3-Kinase regulatory subunit isoforms were immunoprecipitated from cell lysates using an anti-FLAG antibody and the co-immunoprecipitation of ACK and/or dACK was assessed using an anti-HA antibody (top panels). Results are representative of at least three independent experiments.

### ACK phosphorylates p85α, p85β, p55α and p50α at a conserved tyrosine within the iSH2 region

After confirming all five PI3-Kinase regulatory subunits as ACK binding partners, we next examined whether they were also ACK substrates. Several potential phosphotyrosine sites have been identified in the iSH2-cSH2 ACK-interacting region: one of these, Tyr607 (p85α numbering) within the iSH2, is reported to be a major phosphorylation site on p85α for the insulin receptor (22). Using an anti-p85 pTyr607 antibody, we found that p85α, p55α and p85β were phosphorylated on Tyr607, or the equivalent tyrosine residue, when they were co-expressed with ACK but not with dACK (Figure 3A, B and D). Phosphorylation at this site in p50α (Tyr307) could not be detected, however, immunoblotting with a pan anti-pTyr antibody indicted that p50α did show increased tyrosine phosphorylation in the presence of ACK but not dACK (Figure 3C).

**Figure 3:**
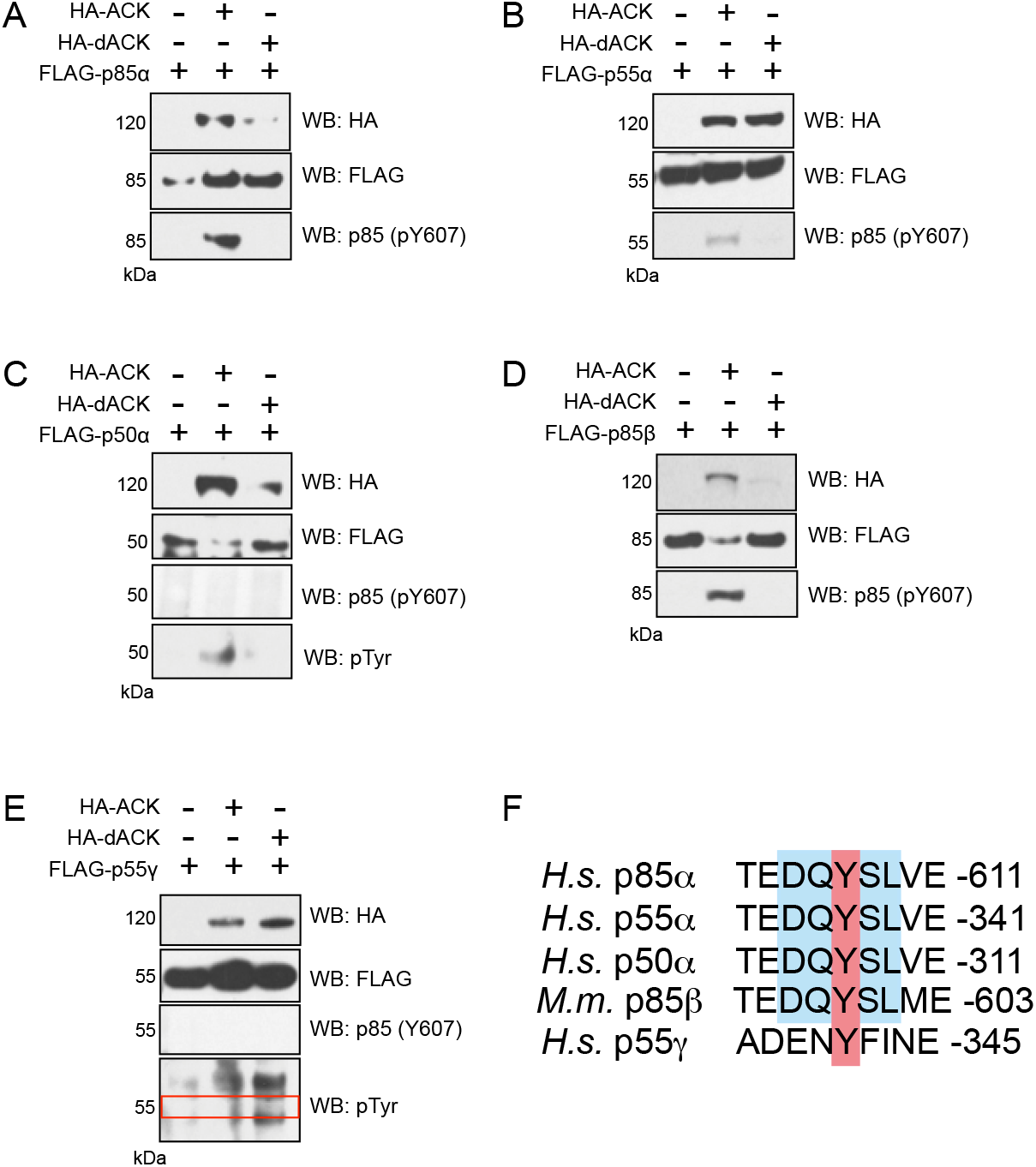
Phosphorylation of full-length FLAG-tagged regulatory subunit isoforms by full-length HA-ACK in HEK293T cells. (A-E) HEK293T cells were harvested 40 hours after transfection with FLAG-tagged regulatory subunit isoforms in the presence and absence of HA-ACK/HA-dACK. Phosphorylation of FLAG-tagged regulatory subunit isoforms was assessed by western blotting of cell lysates using an antibody raised against the p85 sequence (D-Q-Y_(p)_-S-L) and/or a pan anti-pTyr antibody. The red box in (E) denotes the region where a signal for pTyr p55γ would be expected. Results are representative of at least three independent experiments. (F) Amino acid sequence alignment surrounding Tyr607 or equivalent (red) in all p85 isoforms. Sequence recognized by the anti-p85 (pTyr607) antibody is shown in blue.

It was not possible to determine whether p55γ was phosphorylated on Tyr341 (equivalent to Tyr607 in p85α) when co-expressed with ACK using the anti-p85 pTyr607 antibody, as p55γ does not possess the motif (D-Q-Y_(p)_-S-L) recognized by this antibody (Figure 3F). However the change in sequence context around this Tyr in p55γ also implied that it was unlikely to be phosphorylated by ACK. Immunoblotting with a pan anti-pTyr antibody was used to assess whether p55γ showed increased tyrosine phosphorylation when co-expressed with ACK in cells. The signal from the pan anti-pTyr antibody at 55 kDa was no stronger in the lysate co-expressing p55γ and ACK than lysate co-expressing p55γ with dACK (Figure 3E), suggesting that p55γ is not a substrate for ACK.

To confirm that phosphorylation on Tyr607 by ACK was direct, *in vitro* kinase assays were performed using a purified, recombinant, active fragment of ACK. This was used to phosphorylate full-length regulatory subunit isoforms purified from *E. coli*. Incubation of GST-tagged p85α, p85β, p55α and p50α with ACK resulted in their direct phosphorylation on Tyr607 (or equivalent), as assessed by immunoblotting (Figure 4A-D).

**Figure 4:**
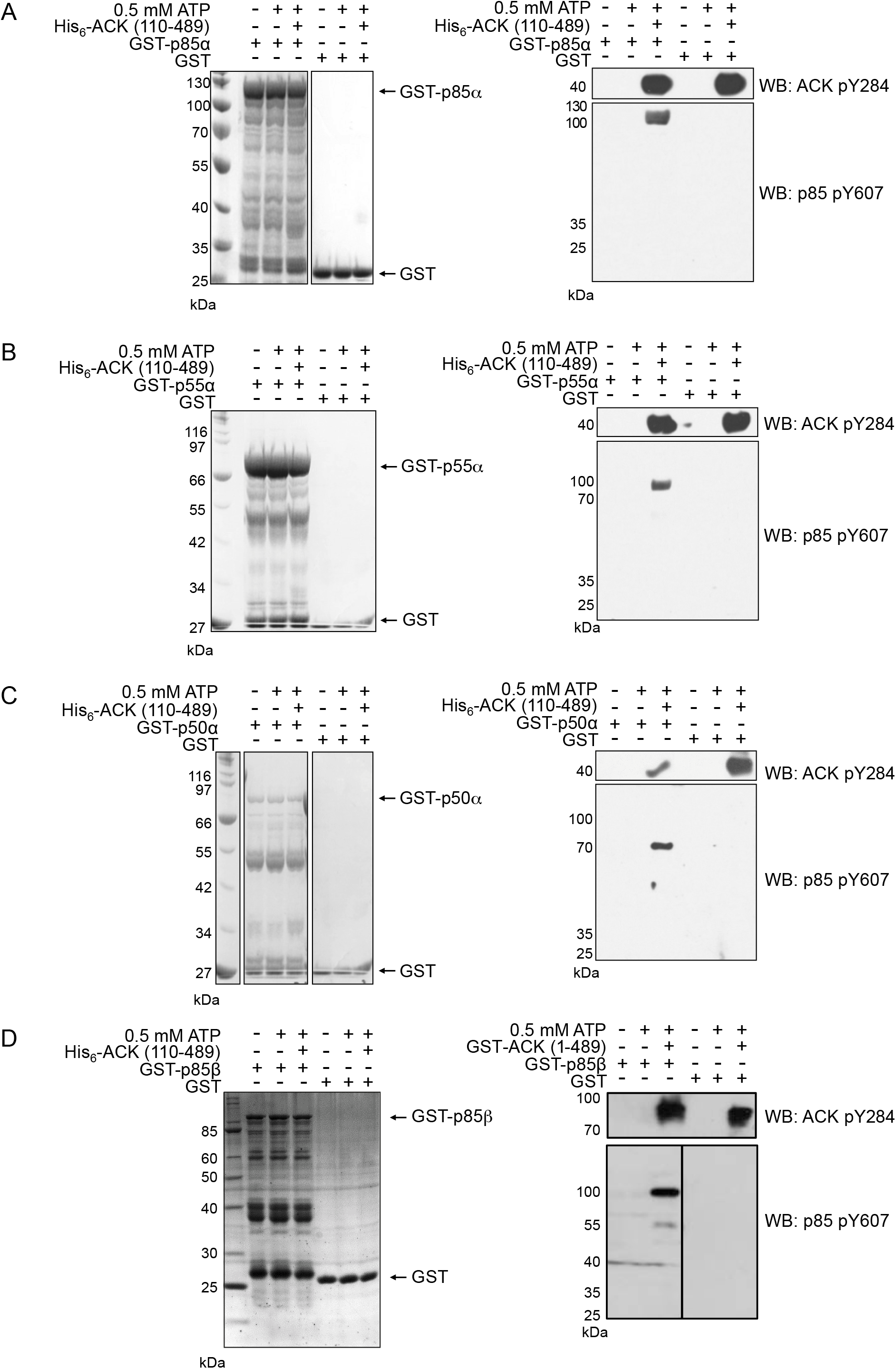
Phosphorylation of the PI3-Kinase regulatory subunits *in vitro*. Full-length GST-tagged regulatory subunits or GST control were incubated at 30 °C for 30 minutes with His_6_-ACK (110-489) or GST-ACK (1-489). Reactions were analysed by SDS-PAGE and Coomassie staining (left panels) and western blotting using anti-p85 (pTyr607) and anti-ACK (pTyr284) antibodies (right panels). Results are representative of at least three independent experiments. (A) p85α is phosphorylated on Tyr607 by purified His_6_-ACK110-189, (B) p55α is phosphorylated on Tyr337, (C) p50α is phosphorylated on Tyr307 and (D) p85β is phosphorylated on Tyr599 by purified GST-ACK (1-489) *in vitro*.

To investigate whether ACK phosphorylates the regulatory subunits at additional tyrosine residues, the *in vitro* kinase assay using GST-p50α and ACK was repeated, resolved by gel electrophoresis and the phosphorylation site(s) mapped by mass spectrometry. Comparison of the mass spectra of samples with and without ACK detected phosphorylation of Tyr307 p50α (equivalent to Tyr607 in p85α) only when ACK was present (Figure S1). No other tyrosine phosphorylation sites were identified, indicating that ACK phosphorylates p50α only at Tyr307 *in vitro*. To confirm that p55γ was not an ACK substrate, His_6_-MBP-p55γ was incubated with ACK and phosphorylation sites were mapped by LC-MS/MS. No increase in p55γ tyrosine phosphorylation was detected following incubation with ACK, corroborating the results obtained in HEK293T cells.

### Phosphorylation of p85β by ACK promotes cell proliferation

Tyr607 (or equivalent) is located within the inter-SH2 region of the regulatory subunits (Figure 1B). The inter-SH2 region is known to provide important regulatory contacts with the PI3-Kinase catalytic subunit (23) and insulin receptor activation results in phosphorylation at this site *in vivo* (22). We hypothesized that phosphorylation of the regulatory subunits by ACK within this functionally significant region would lead to an increase in PI3-Kinase catalytic activity, which could in part mediate the increase in cell proliferation that has been observed following overexpression of constitutively active ACK (caACK) in cells in culture (12). To investigate this, HEK293T cell lines were generated which stably expressed wild type (wt) ACK, caACK, wt p85β or a non-phosphorylatable p85β mutant, Y599F (equivalent to Tyr607 in p85α). Several lines were analyzed for equivalent protein expression levels and then tested for comparative rates of proliferation. Proliferation assays confirmed that the introduction of either wt or caACK into HEK293T cells conferred a proliferative advantage (Figure 5A). Cells expressing wt p85β grew at a slower rate than control cells (Figure 5B) demonstrating that p85β inhibits proliferation. This is in agreement with previous studies that indicate increased levels of p85 negatively regulate PI3-Kinase activity (24, 25). Cells expressing Y599F p85β however, proliferated at an even slower rate suggesting that the inhibitory effect of p85β is counteracted by phosphorylation at Tyr599. The effects of the ACK specific inhibitor AIM-100 (26) were also tested. Cells expressing wt p85β grew at a reduced rate in the presence of AIM-100 (Figure 5B), suggesting that phosphorylation at Tyr599 by endogenous ACK counteracts some of the anti-proliferative effects of p85β, while no effect was seen for cells expressing Y599F p85β in the presence of AIM-100. Overall these data suggest that ACK stimulates cell proliferation by phosphorylating p85β at Tyr599.

**Figure 5:**
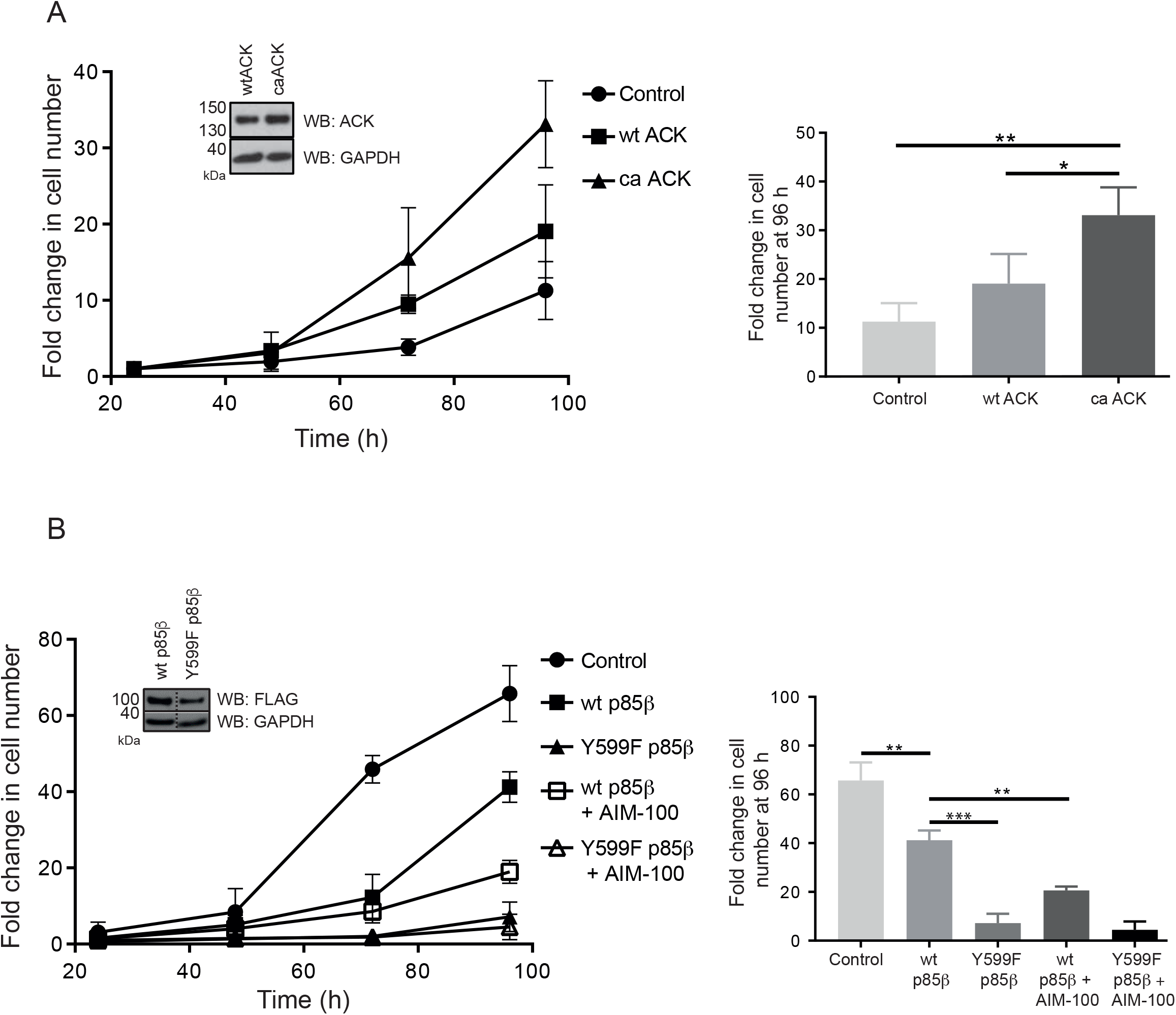
Proliferation of HEK293T cells expressing ACK and p85β variants. (A) Proliferation of HEK293T cells stably expressing wt ACK and caACK. HEK293T cells containing empty vector or stably expressing HA-wt ACK or caACK were seeded at a density of 3 × 10^4^ cells per well. Cells were plated in replicates of 5 and live cell counts were obtained at the times indicated. Data from three independent experiments is shown. Error bars represent standard deviation. Inset shows the relative levels of ACK in the wt and caACK cell lines by western blot analysis. Data from the 96 h time point is also shown in a bar chart on the right hand side. Group significance testing was performed by T-test: *, p < 0.05; **, p < 0.01. B) Proliferation of HEK293T cells stably expressing wt p85β and Y599F p85β. HEK293T cells stably expressing inducible FLAG-tagged wt p85β or p85β Y599F were seeded at a density of 3 × 10^4^ cells per well and incubated at 37 °C for 16 hours. Cells were then treated with 1µg/mL doxycycline and incubated at 37 °C for the times indicated before live cell counts were taken. Certain cell lines were treated with the ACK inhibitor AIM-100 (8 μM) as indicated. Data from three independent experiments is shown. Error bars represent standard deviation. Inset shows the relative levels of p85 in the wt and Y599F cell lines by western blot analysis. Data from the 96 h time point is also shown in a bar chart on the right hand side. Group significance testing was performed by T-test: *, p < 0.05; **, p < 0.01, ***, p < 0.001.

### ACK suppresses increases in PIP_3_ following EGFR activation

We rationalized that phosphorylation of the regulatory subunits might increase the catalytic activity of PI3-Kinase by modulating the interaction between the regulatory and catalytic subunits. Mass spectrometry was used to quantify levels of PIP_3_ directly in cells that had been transfected with ACK. To quantify total cellular PIP_3_, cellular phosphoinositides were extracted from HEK293T cells and derivatized (19). Extracts were then analysed by high-performance liquid chromatography-mass spectrometry. To assess whether ACK affects basal and/or EGF-stimulated PIP_3_ levels in HEK293T cells, cells were transfected with constructs expressing wtACK or caACK. After 24 hours, cells were serum starved and then treated with EGF to stimulate PIP_3_ production. Whilst control (GFP) transfected cells showed an increase in PIP_3_ following EGF stimulation, this was not observed in cells transfected with wtACK (Figure 6), suggesting that ACK actually suppresses the PI3-Kinase response to EGF stimulation. Consistent with this, expression of caACK decreased levels of PIP_3_ following EGF treatment (Figure 6). These data suggest that ACK either suppresses the generation of PIP_3_ or promotes PIP_3_ clearance, effects that are likely to be dependent on ACK kinase activity. Moreover the data suggest that stimulation of cell proliferation by phosphorylation of p85 at Tyr607 by ACK is not mediated through an increase in PI3-Kinase lipid kinase activity but by an alternative mechanism.

**Figure 6:**
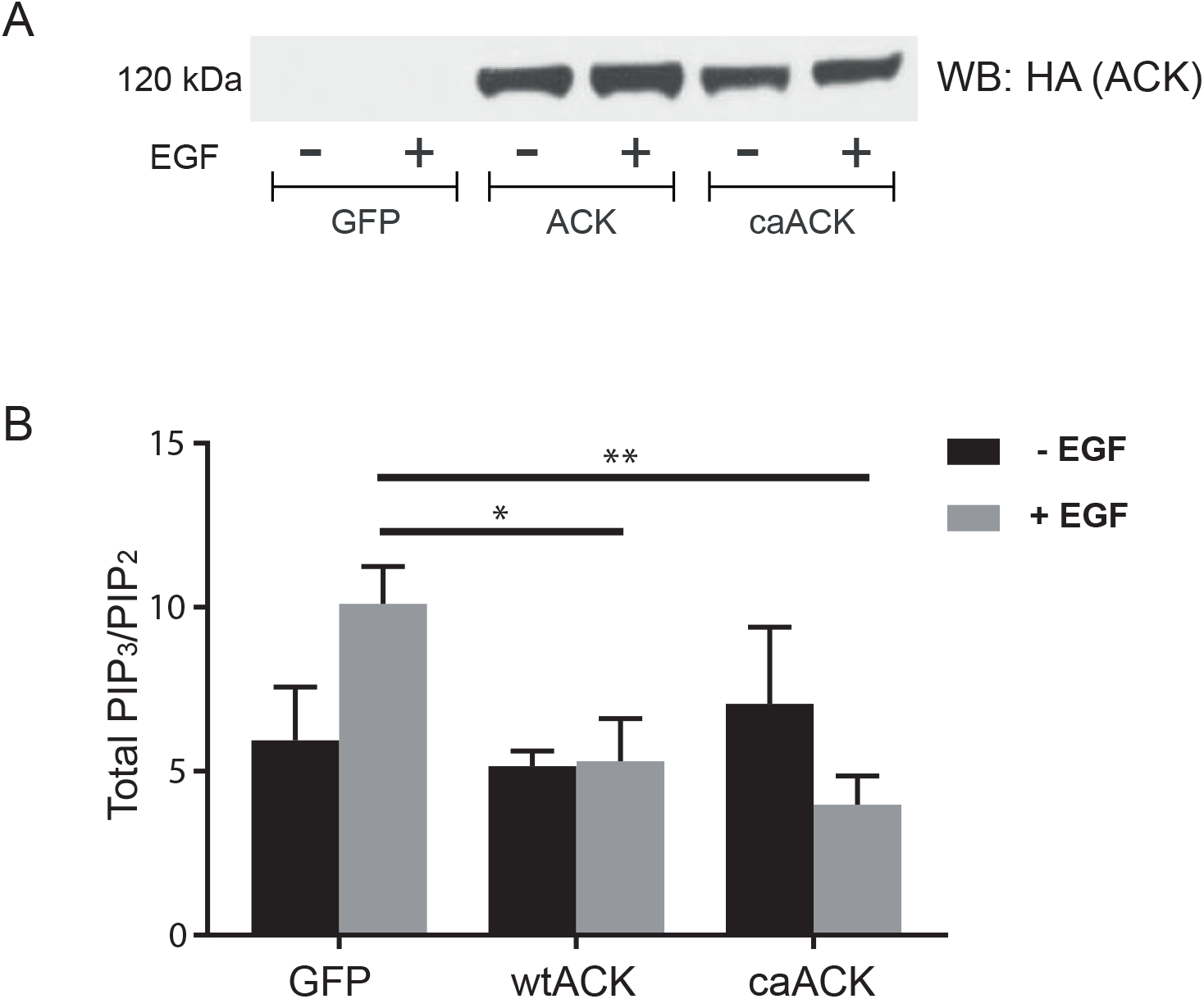
Effect of exogenous wtACK and caACK expression on PIP_3_ levels in serum-starved HEK293T cells. Cells were transfected with GFP, HA-wtACK or HA-caACK for 24 hours. Cells were then grown in 0% FBS DMEM for 16 hours before stimulation with EGF. (A) Whole cell lysates from one experimental replicate were normalised by Bradford assay and 80 μg of each lysate was analysed by Western blotting using an anti-HA antibody to confirm similar levels of expression of HA-ACK and HA-caACK. (B) PIP_3_ and PIP_2_ were extracted and quantified using HPLC-MS. PIP_3_ levels are expressed as a ratio of the mean total cellular PIP_3_ to mean total cellular PIP_2_ (error bars show the standard deviation from n=3 technical repeats). Responses were normalized for cell input using a 16:0/17:0-PIP_3_ internal standard. Data is representative of trends observed in three independent experiments.

### Stabilization of p85 by ACK

There is considerable evidence that increased levels of cellular p85α and p85β result in inhibition of PI3-Kinase activity (27, 28). The mechanism through which this inhibition is achieved is not clear, although it has been suggested that p110-free p85 could compete with PI3-Kinase heterodimers for binding to phosphotyrosine motifs on activated RTKs (28). To investigate whether ACK increases the levels of p85α/β in HEK293T cells by stabilising the p85 protein, levels of endogenous p85α/β in the presence and absence of transfected ACK were compared following treatment of cells with cycloheximide (Figure 7A). In the absence of exogenous ACK expression, endogenous p85α/β levels are significantly reduced after cycloheximide treatment. This effect is not seen in cells that have been transfected with ACK, suggesting that the level of p85 in the cell is, at least in part, regulated by ACK-mediated stabilization.

**Figure 7:**
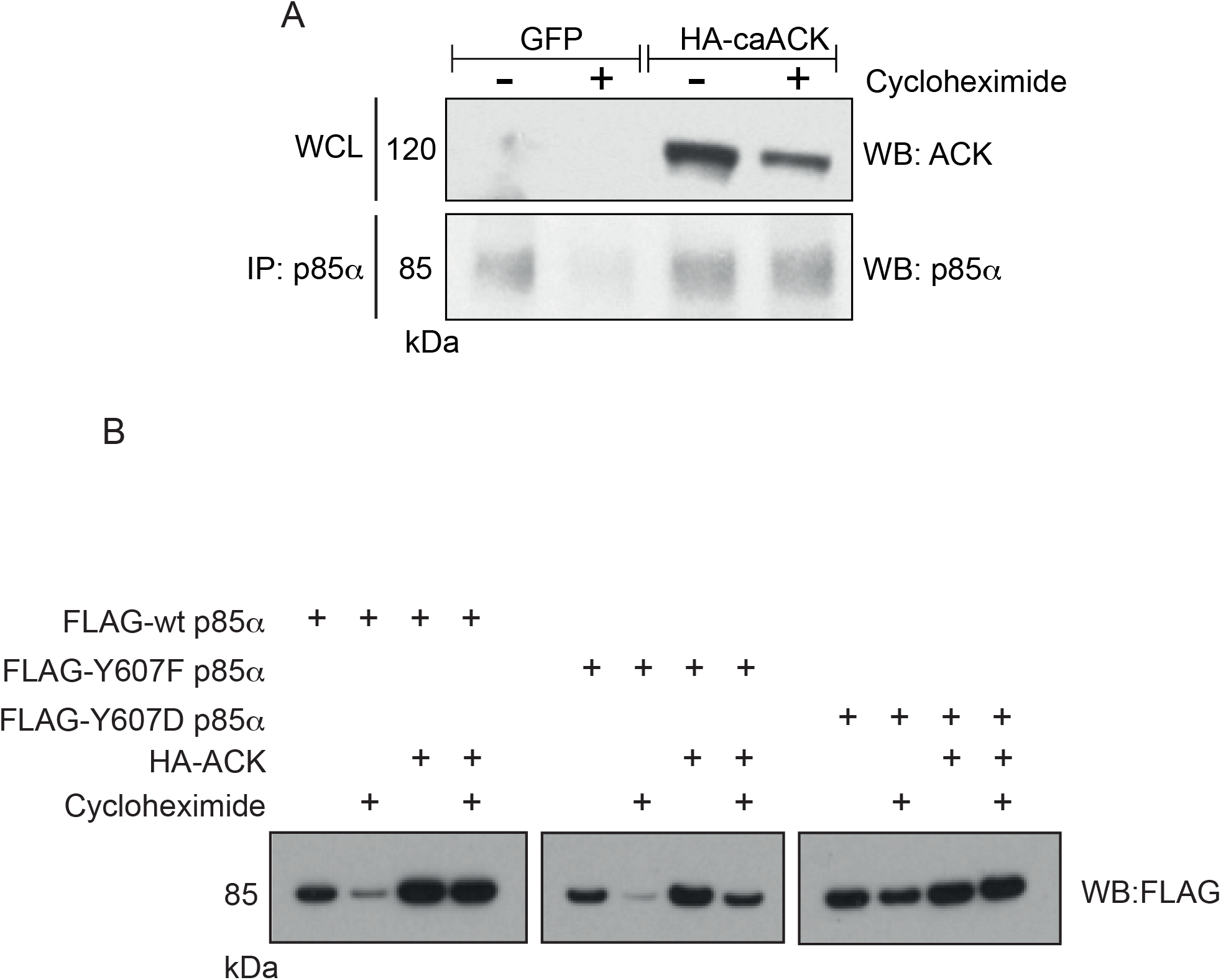
Stabilization of p85α by ACK *in vivo*. (A) HEK293T cells were transfected with either HA-caACK or HA-GFP and then treated with cycloheximide (+) or DMSO (-). Cell lysates were immunoprecipitated using an anti-p85α antibody. Western blotting with an anti-ACK antibody was used to assess the expression of HA-caACK in whole cell lysates (top panel) and an anti-p85α antibody was used to assess the amount of immunoprecipitated endogenous p85α. (B) HEK293T cells were transfected with FLAG-wt p85α, FLAG-Y607F p85α or FLAG-Y607D p85α, +/− HA-ACK and then treated with cycloheximide (+) or DMSO (-). Cells were lysed, supernatants quantified for protein levels and equal amounts of lysate protein extracts were immunoprecipitated using an anti-p85 antibody. Levels of FLAG-p85α variants were assessed by western blotting using an anti-FLAG antibody. All data is representative of at least three independent experiments.

To investigate whether the mechanism underpinning stabilization of p85 by ACK involves phosphorylation at Tyr607, HEK293T cells were transfected with wt p85α, p85α Y607D, a phosphomimic, or p85α Y607F, which cannot undergo phosphorylation at this site. Although substitution of tyrosine with aspartate is not an ideal phosphotyrosine mimic, it can be used effectively in some cases and this approach has been utilized successfully, previously, for analysing p85 SH2 domain binding to phosphotyrosine sequences (29). The levels of wt p85α dropped considerably after exposure to cycloheximide but were entirely retained in the presence of ACK (Figure 7B). It is also notable that the level of wt p85α in the absence of cycloheximide is higher in the presence of ACK. Y607F p85α was degraded to very low levels in the presence of cycloheximide and although levels were higher in the presence of ACK, a clear reduction was observed after treatment with cycloheximide (Figure 7B). In contrast, levels of Y607D p85α decreased only slightly with cycloheximide treatment and, similar to wt p85α, the levels are higher in the presence of ACK (Figure 7B). These data indicate that ACK-dependent phosphorylation of p85α Tyr607 stabilizes p85α and maintains elevated protein levels.

### The mechanism underpinning the stabilization of p85α by phosphorylation of Tyr607

The ubiquitin-proteasome pathway regulates the stability of the p85α protein. The tumour suppressor p42 has been reported to promote degradation of p85α by targeting its iSH2 domain (30). We have shown that the level of p85α in the cell is in part regulated by ACK and that this regulation is achieved through protection of p85α from degradation. Intriguingly, the site of ACK phosphorylation, Tyr607, is located within the iSH2 domain of p85α, which is flanked by two SH2 domains (Figure 1B). We hypothesized that p85 stabilization might be due to an intramolecular interaction between one of these SH2 domains and phosphoTyr607, which would result in protection of the iSH2 region from the ubiquitin-proteasome machinery.

To test this hypothesis we expressed and purified each SH2 domain from p85α and p85β separately in *E. coli.* We then tested their ability to interact with peptides containing the target tyrosine and surrounding sequence from p85α and p85β, using a fluorescence polarization assay to monitor binding. The nSH2 domains of both p85α and p85β interacted with a p85α pY607 peptide with low micromolar affinity but bound with high affinity (low nanomolar) to a p85β pY607 peptide (Figure 8 and Table 1). No interaction was seen with the non-phosphorylated versions of the same sequences (Figure 8A and B) and no binding was observed between either peptide and either of the cSH2 domains (Figure 8C and D). To confirm that the cSH2 domains were functional, binding was tested to a known target peptide from CD28 (Figure 8C and D). Binding affinities similar to those previously reported were observed, indicating that both cSH2 domains were capable of binding phosphopeptides (31). Thus pTyr607 supports an interaction with p85α and p85β nSH2 domains, with higher affinity for the p85β pY607 peptide.

**Table 1:** Equilibrium binding constants for p85 SH2 domains and phosphorylated peptides

| $K_d$ (nM)<sup>a</sup> | | | |
| --- | --- | --- | --- |
| | pYSLV ($\alpha$ peptide) | pYSLV ($\beta$ peptide) | pCD28 peptide |
| p85α nSH2 | 2400 ± 315 | 82 ± 16 | - |
| p85β nSH2 | 9000 ± 282 | 52 ± 8 | - |
| p85α cSH2 | 84000 | 656 ± 424 | 43 ± 10 |
| p85β cSH2 | ND<sup>b</sup> | ND | 100 ± 27 |
| p85α nSH2 R358M | ND | 2900 ± 1600 | - |
| p85β nSH2 R349M | ND | ?? | - |

<sup>a</sup> Standard error where n=3
<sup>b</sup> ND (no binding) denotes data that could not be fitted to the binding isotherm

**Figure 8:**
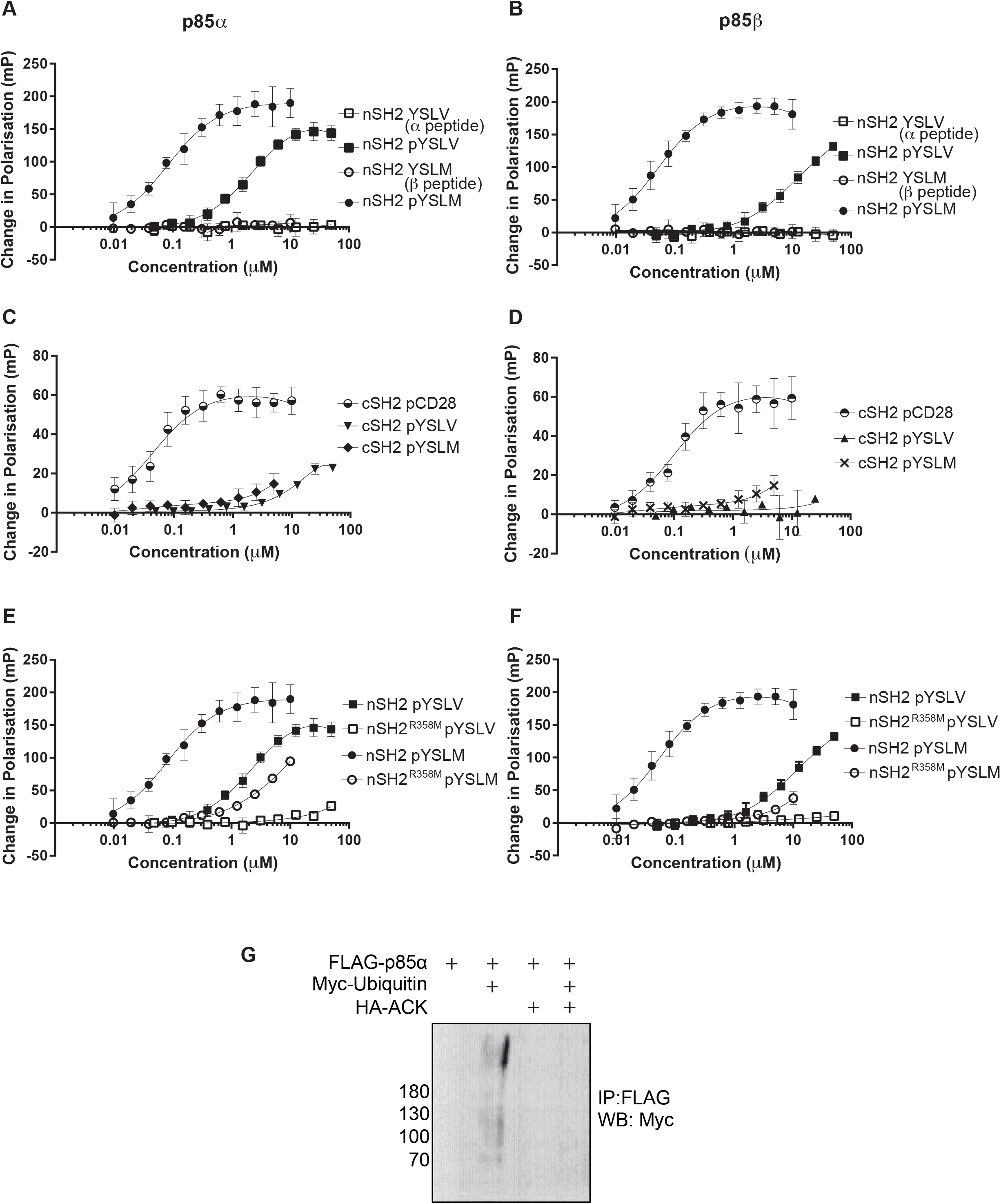
Binding of the p85 SH2 domains to phosphopeptides. Purified nSH2 and cSH2 domains from p85α and p85β were analysed for binding to synthetic phosphopeptides based on the sequences surrounding Tyr607 in p85α and p85β, in fluorescence polarization assays. Data from three independent experiments is shown with error bars represent standard deviation. (A) Binding of the nSH2 domain of p85α to phosphorylated and non-phosphorylated peptides based on p85α and p85β. (B) Binding of the nSH2 domain of p85β to phosphorylated and non-phosphorylated peptides based on p85α and p85β. (C) Binding of the cSH2 domain of p85α to phosphorylated peptides based on p85α and p85β, and a control peptide from CD28. (D) Binding of the cSH2 domain of p85β to phosphorylated peptides based on p85α and p85β, and a control peptide from CD28. (E) Binding of the wt nSH2 domain of p85α and the R358M variant to phosphorylated peptides based on p85α and p85β. (F) Binding of the wt nSH2 domain of p85β and the R358M variant to phosphorylated peptides based on p85α and p85β. (G) Ubiquitination of p85α. FLAG-p85 was expressed +/− Myc-Ubiquitin and +/− HA-ACK in HEK 293T cells. Cell lysates were immunoprecipitated with anti-FLAG antibody and then analysed by Western blotting with an anti-Myc antibody.

We further tested the binding mode of the p85 nSH2 domains. An arginine residue is evolutionarily conserved amongst SH2 domains and forms a hydrogen bond with the phosphate group of the phosphotyrosine residue on binding targets (32). Mutation of this arginine to methionine in p85α completely abolishes phosphopeptide binding without disrupting the characteristic SH2 fold (31). Site-directed mutagenesis was used to generate SH2 mutants p85α R358M and p85β R349M and the purified domains were analysed in binding assays. These nSH2 null mutations reduced binding to the phosphopeptide sequence by several orders of magnitude, indicating that the pTyr607-wildtype nSH2 binding mechanism is likely to be similar to other canonical p85 SH2-peptide interactions (Figure 8E and F).

As targeting of the iSH2 domain in p85 has been shown to induce degradation by the ubiquitin pathway (30), we speculated that phosphorylation at Tyr607 and subsequent interaction with the nSH2 domain would interfere with this process. HEK293T cells were transfected with FLAG-tagged p85α, Myc-tagged Ubiquitin and HA-tagged ACK and then treated with MG132, a proteasomal inhibitor, or DMSO as a control. The cell lysates were immunoprecipitated with an antibody specific for the FLAG tag and ubiquitinated p85α detected with an anti-Myc antibody. Polyubiquitinated p85α protein was detected in the MG132 treated samples as expected (Figure 8G). Co-expression of ACK however, completely abolished ubiquitination of p85α. Taken together these results indicate that phosphorylation of p85α Tyr607 by ACK leads to an nSH2-phosphotyrosine interaction that blocks access to the ubiquitination machinery and thus regulates p85α protein stability.

### ACK interacts with p85α in nuclear-enriched cell fractions

We have demonstrated that ACK phosphorylates the PI3-Kinase regulatory subunits on Tyr607 and that this promotes cell proliferation through a mechanism that does not involve an increase in lipid kinase activity. We hypothesized that phosphorylation at Tyr607 by ACK may instead promote cell proliferation by shunting PI3-Kinase or its subunits into alternative pathways independent of lipid kinase activity. We also know that phosphorylation of p85 inhibits ubiquitin-dependent degradation likely resulting in an increase in p110-free p85 subunits. The documented p110-independent functions of p85 predominate in the nucleus (33, 34).

To investigate whether ACK promotes cell proliferation by modulating these kinase-independent roles of PI3-Kinase or its subunits in the nucleus, we first determined the subcellular localization of ACK and the PI3-Kinase regulatory subunits and identified their site of interaction. Whilst PI3-Kinase and ACK are primarily activated by RTKs at the plasma membrane, nuclear translocation of ACK (14, 35) and the p85α regulatory subunit (33, 36) have been documented. To investigate whether co-expression of ACK and p85α altered the subcellular localization of either of the two proteins, HEK293T cells were transiently transfected with ACK and p85α expression constructs and then separated into cytoplasmic and nuclear-enriched fractions.

ACK was found in both the cytoplasmic and nuclear-enriched fractions and this subcellular localization was not affected by co-expression with p85α (Figure 9A). This suggests that ACK is able to shuttle between the nucleus and cytoplasm via a mechanism that is not regulated by its interaction with p85α. p85α was also detected in both cellular fractions but was predominantly (~90%) in the nuclear-enriched fraction when expressed alone (Figure 9A). Co-expression with ACK increased the total amount of p85, as seen previously, and resulted in a near equal distribution of p85 between nucleus and cytoplasm.

**Figure 9:**
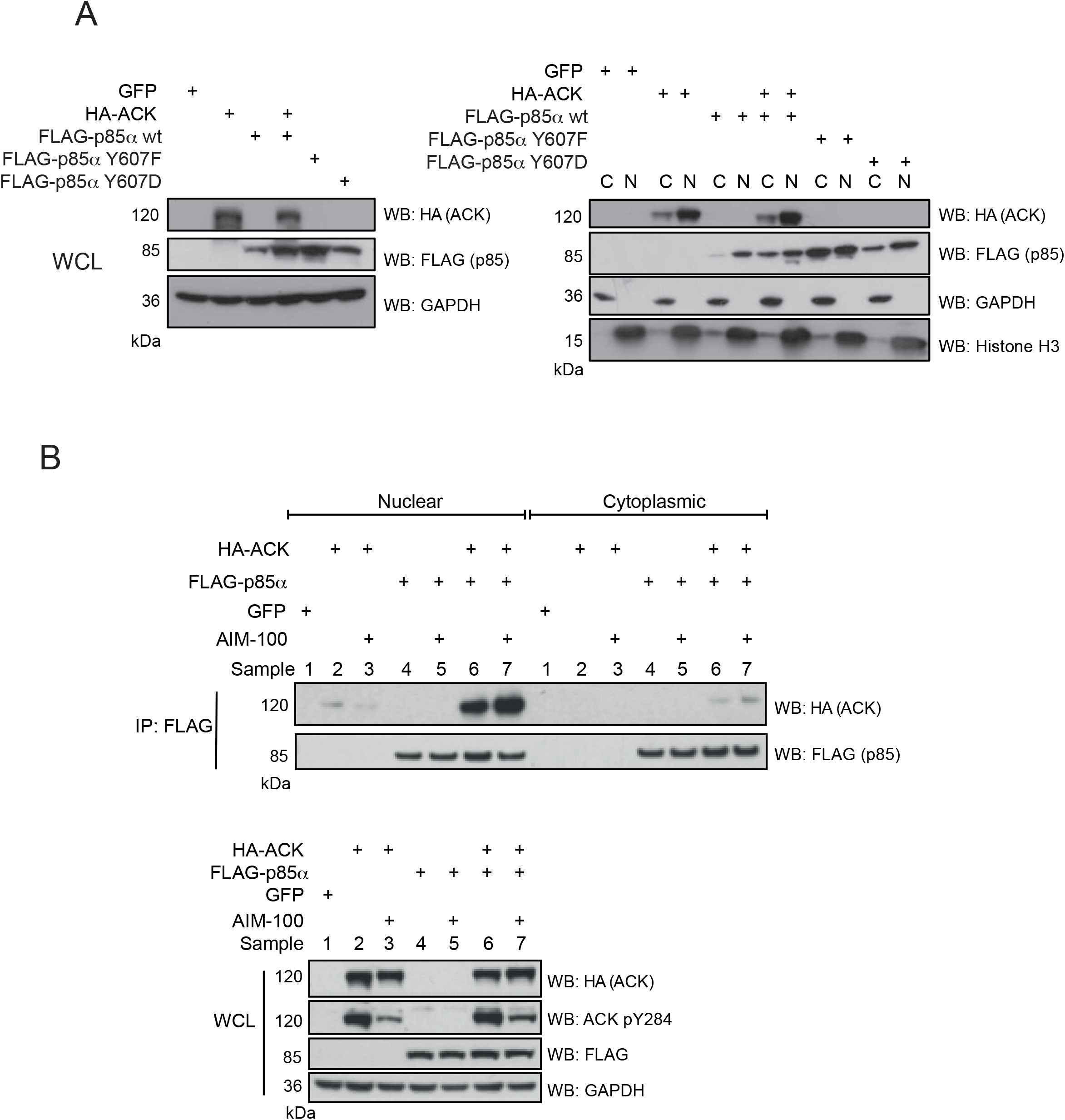
Subcellular localization of ACK, p85α and the ACK-p85α complex. (A) ACK and wt p85α were expressed alone and together in HEK293T cells. Y607D p85α and Y607F p85α were expressed alone. Samples of whole cell lysates (WCL) were analysed by western blotting using the antibodies indicated and are shown in the left hand panels. Cells were lysed and separated into cytoplasmic and nuclear-enriched fractions. Fractions were analysed by western blotting using the antibodies indicated (right hand panels). Fractionation was judged using anti-GAPDH to identify cytoplasmic extracts and anti-Histone H3 to identify nuclear-enriched fractions. The amount of p85α variants found in the cytoplasmic and nuclear-enriched fractions were analysed by densitometry using Image J. The total amount of p85 was normalized to 100 and the fractions expressed as a percentage of total. wt p85α expressed alone was distributed 7:93 cytoplasmic (C):nuclear (N); p85α co-expressed with ACK, 46:54; p85α Y607F 50:50 and p85α Y607D, 30:70. (B) Top panels: FLAG-p85α was immunoprecipitated from nuclear-enriched and cytoplasmic fractions using an anti-FLAG antibody. Immunoprecipitation of FLAG-p85α and co-immunoprecipitation of HA-ACK was assessed by western blotting using anti-FLAG and anti-HA antibodies. Bottom panels: Expression of FLAG-p85α and HA-ACK along with active ACK levels was confirmed by western blotting of whole cell lysate (WCL) samples collected prior to cell fractionation. All data is representative of at least three independent experiments.

To determine the subcellular location of ACK-p85α complexes, cellular fractions were immunoprecipitated with an anti-FLAG antibody. The interaction between p85α and ACK was detected almost exclusively in nuclear-enriched fractions (Figure 9B) suggesting that phosphorylation of p85 by ACK occurs in the nucleus. Consistent with this, p85 pY607 was almost entirely located in the nuclear fractions (Figure S2).

To investigate whether phosphorylation of p85α on Tyr607 influences its cellular location, the subcellular localisation of the p85α mutants Y607D and Y607F was determined (Figure 9A). Whereas p85α Y607F was equally distributed between the cytoplasmic and the nuclear-enriched fractions, a higher percentage of p85 Y607D (~70%) was nuclear suggesting that Y607 phosphorylation influences the subcellular localization of p85.

### The pTyr607-nSH2 interaction occurs *in trans* and promotes formation of novel p85 dimers

The interaction between the nSH2 and pTyr607 of the regulatory subunits could occur theoretically either *in cis* in a monomeric protein, or *in trans* to form a dimer. Either conformation would protect the iSH2 domain from targeting by E3 ligases. To investigate whether the pTyr607-nSH2 interaction could support dimer formation we co-expressed either FLAG-p50α with V5-p85β or FLAG-p85α with V5-p85β. We chose to study p85β as peptides from this isoform had the highest affinity binding to the nSH2 domains tested. To distinguish between dimers formed through interactions located in the N-terminal or C-terminal half of the molecules we also utilized the p50α isoform, which lacks the N-terminal SH3, polyproline and RhoGAP/BH regions.

Proteins were co-expressed in HEK293T cells cultured in the presence of either 10% serum or in low serum (0.05%) followed by stimulation with EGF or insulin. p50α interacted with p85β under all conditions, although lower levels of complex were seen in low serum. Increased p50α-p85β complex formation was observed in high serum or in response to EGF or insulin stimulation (Figure 10A), suggesting that dimerization is induced by growth factor signalling. Dimerization in this case must be occurring solely via C-terminal interactions. p85α was also seen to interact with p85β (Figure 10B). Again, lower levels of complex were seen in low serum with an increase in 10% serum or following insulin or EGF stimulation The increase in this case was more modest, potentially reflecting the presence of additional N-terminally mediated dimers (Figure 10B). To investigate whether the interactions observed were mediated by the pTyr607-nSH2 interaction we determined the interaction with a p50α phosphomimic mutation (Y307D, equivalent to Y607D in p85) (Figure 10C). Whereas in low serum conditions, wt p50α -p85β interaction was observed at low levels, as previously observed, a significantly increased level of p50α Y307D-p85β dimer was seen, suggesting that phosphorylation of Tyr307 underpins this interaction.

**Figure 10:**
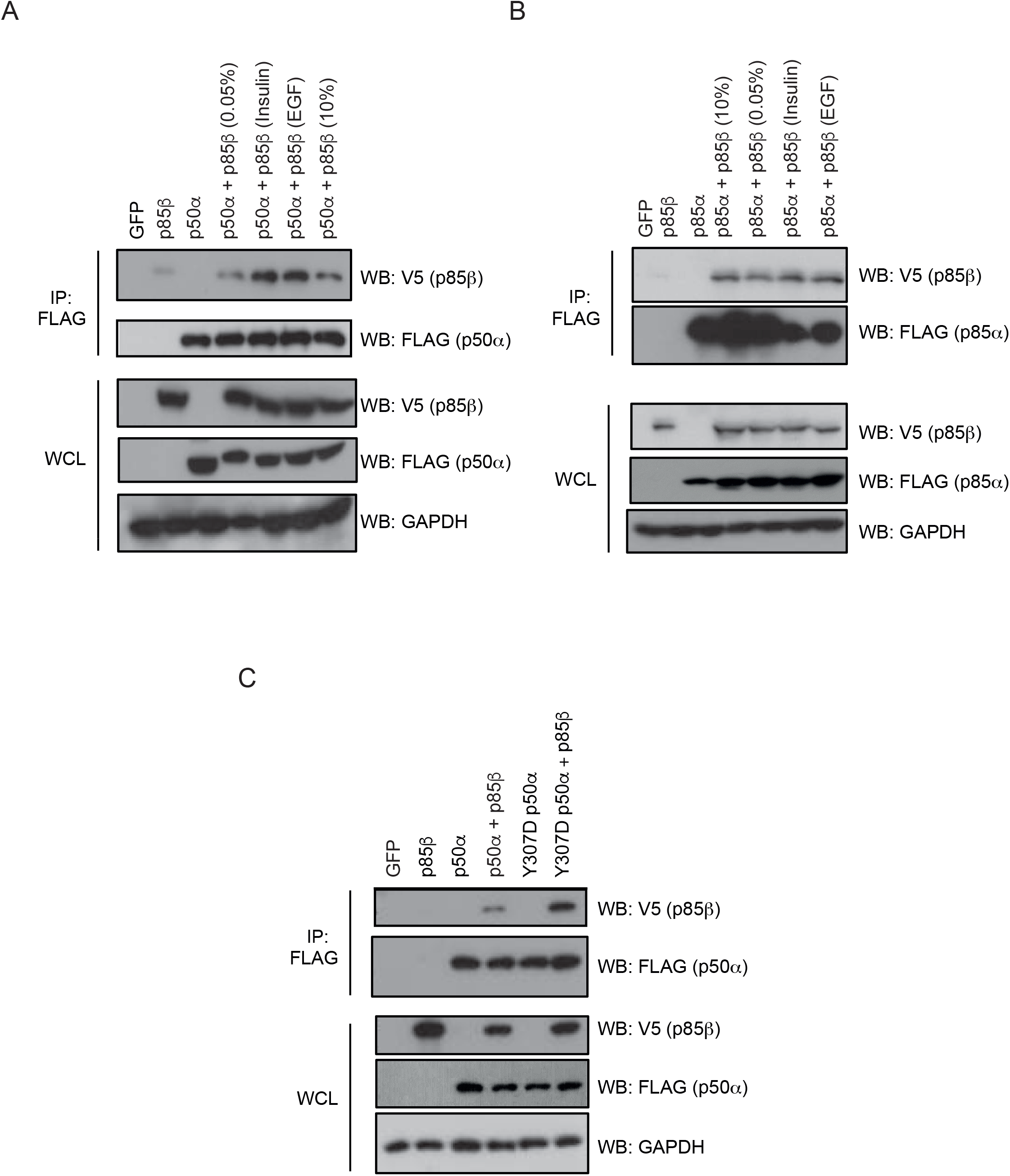
Dimerization of the regulatory subunits. (A) wt p85β and wt p50α were expressed alone or co-expressed in HEK293T cells. Samples of whole cell lysates (WCL) were analysed by western blotting using the antibodies indicated and are shown in lower panels. p50α was immunoprecipitated from cell lysates using an anti-FLAG antibody and the co-immunoprecipitation of p85β was assessed using an anti-V5 antibody (top panels). Results are representative of at least three independent experiments. (B) wt p85β and wt p85α were expressed alone or co-expressed in HEK293T cells. Samples of whole cell lysates (WCL) were analysed by western blotting using the antibodies indicated and are shown in lower panels. p85α was immunoprecipitated from cell lysates using an anti-FLAG antibody and the co-immunoprecipitation of p85β was assessed using an anti-V5 antibody (top panels). Results are representative of at least three independent experiments. (C) wt p85β, wt p50α and Y307D p50α were expressed alone or co-expressed in HEK293T cells. Samples of whole cell lysates (WCL) were analysed by western blotting using the antibodies indicated and are shown in lower panels. p50α variants were immunoprecipitated from cell lysates using an anti-FLAG antibody and the co-immunoprecipitation of p85β was assessed using an anti-V5 antibody (top panels). Results are representative of at least three independent experiments.

These data demonstrate that the C-terminal regions, and specifically the pTyr607-nSH2 interaction, support formation of a novel dimeric configuration of the regulatory subunits in response to growth signals.

Since both the ACK-p85α interaction and p85 pTyr607 occur exclusively in the nucleus (Figure 9B), we investigated the subcellular location of regulatory subunit dimers. Proteins were co-expressed and the cell fractionated before co-immunoprecipitation. p50α-p85β dimers were found mainly in the nucleus, whereas p85α-p85β dimers were located in both the cytoplasm and the nucleus (Figure 11). We interpret these data to indicate that p50α-p85β dimers, as obligate C-terminal dimers, are mainly found in the nucleus, whereas p85α-p85β, which are likely to be a mixture of both C-terminal and N-terminal dimers, are found in both cellular compartments. The nuclear subset is likely to comprise C-terminal dimers while the cytoplasmic will be in the N-terminal dimer configuration.

**Figure 11:**
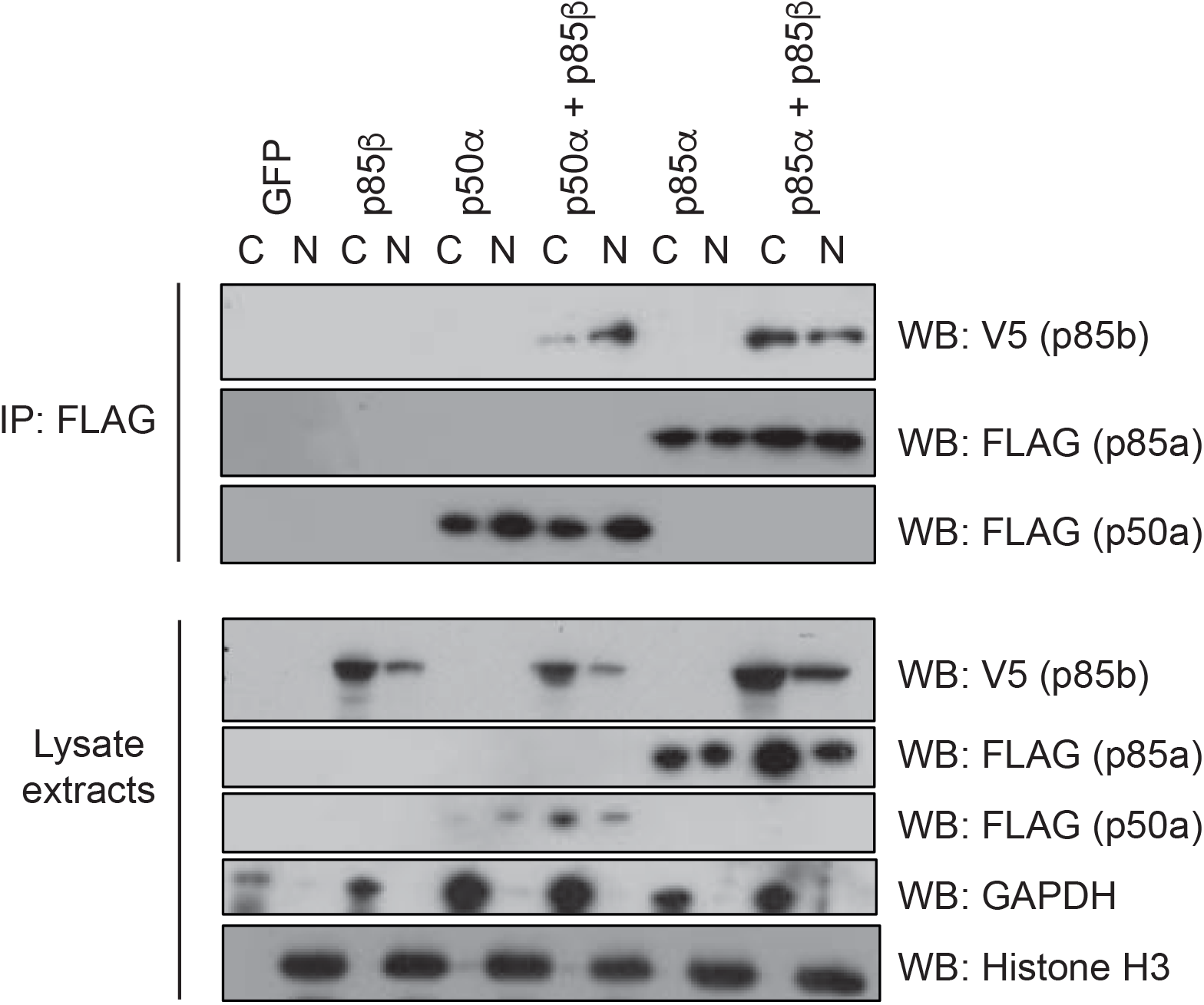
Subcellular localization of the nSH2-pTyr607 regulatory subunit dimers. wt p85β, wt p85α and wt p50α were expressed alone or together in HEK293T cells. Cells were lysed and separated into cytoplasmic and nuclear-enriched fractions. Fractions were analysed by western blotting using the antibodies indicated (lower panels). Fractionation was judged using anti-GAPDH to identify cytoplasmic extracts and anti-Histone H3 to identify nuclear-enriched fractions. p50α or p85α were immunoprecipitated from cell lysates using an anti-FLAG antibody and the co-immunoprecipitation of p85β was assessed using an anti-V5 antibody (top panels). Results are representative of at least three independent experiments.

## Discussion

We have identified the five isoforms of the class Ia PI3-Kinase regulatory subunit as novel ACK binding partners and shown that p85α, p85β, p55α and p50α are also substrates for ACK kinase activity. These isoforms are phosphorylated by ACK at a conserved tyrosine residue (equivalent to Tyr607 in p85α) within the inter-SH2 domain of the protein, a region involved in contacting and regulating the p110 catalytic subunit of PI3-Kinase. We have shown that phosphorylation of the p85β isoform at this site promotes cell proliferation in culture. This increase in cell proliferation is, unexpectedly however, not achieved through a stimulation of PI3-Kinase lipid kinase activity; rather, the increase in PIP_3_ levels that is normally observed following EGF stimulation is suppressed. We have discovered that phosphorylation of p85 at Tyr607 results in stabilization of the protein by protecting it from ubiquitination and subsequent degradation by the proteosome pathway. We propose a mechanism for increased p85 stability mediated by an interaction involving the p85 nSH2 domain and the ACK phosphorylation site (Tyr607 or equivalent). This interaction likely occurs via a binding mode similar to those already described for the p85 nSH2 domain (31), as shown by abrogated binding in the R358M SH2 null mutant. It is likely that the pTyr607-nSH2 interaction masks the iSH2 region, the target site on p85 for the degradative machinery (30). Supporting this, we have shown that ACK protects p85 from polyubiquitination, with ACK expression completely abolishing ubiquitination of p85. Together, our data indicate that ACK, at least in part, regulates the cellular level of p85 by blocking its degradation. Indeed in their paper identifying p42 Ebp1 as the E3 ubiquitin ligase for p85, Ko *et al.* speculate that their results indicate a possible regulatory role for the nSH2 in protecting the iSH2 from the degradative machinery (30). The pTyr607-nSH2 interaction would provide a mechanism for this regulation. We have shown here that this new mechanism of protein stabilization operates in both p85α and p85β and, we assume, will be present for the other two analogous shorter isoforms, p55α and p50α.

We were initially surprised by our observation that active ACK, which is normally pro-proliferative, is associated with a decrease in PIP_3_ generation in HEK293T cells. However the adjacent site on p85, Ser608, has also been identified as a phosphorylation target, in this case for the PI3-Kinase p110 subunit. This intrinsic regulation in PI3-Kinase also decreases the activity of the enzyme, although the mechanism has not been elucidated (37, 38). Tyr607 itself has also been identified previously as a target site that is phosphorylated downstream of the insulin receptor, although the consequence of this for PI3-Kinase lipid kinase activity was not determined (22). The decrease in PIP_3_ levels following phosphorylation of Tyr607 can however potentially be explained by our observation that ACK stabilizes cellular levels of the regulatory subunits and this could manifest mechanistically via two possible routes. We have shown that ACK inhibits degradation of the p85α isoform, which would lead to an excess of regulatory subunits over catalytic subunits. p110-free p85α exists(24) and can form homodimers (39). The formation of both p110-free monomeric p85 and dimeric p85 would be promoted by elevated levels of p85. p110-free p85 is thought to compete with PI3-Kinase heterodimers for binding to pYXXM motifs on activated RTKs. This dampens the activation of PI3-Kinase in response to growth factors and could explain the reduced levels of PIP_3_ we observed following EGF stimulation of cells overexpressing ACK. More recent work has shown that p85α homodimers can also bind to unphosphorylated PTEN at the plasma membrane and stimulate its catalytic activity (40). It is therefore also possible that the ACK-driven increase in p110-free p85α, leads to an increase in p85 homodimers, which promotes PTEN activity, leading to decreased PIP_3_ levels.

It is also apparent from our work that the interaction between pTyr607 and the nSH2 domain in the regulatory subunits underpins a novel mechanism of dimerization for these proteins. Using the wealth of structural data available for p85α, we constructed a model of how these dimers could form (Figure 12). Structural data is not available currently for p85α residues 599-611, so p85α residues 604-611 were modelled into the structure based on the known binding site of a phosphopeptide on the p85α nSH2 domain. Using the available structural data, an intramolecular monomeric interaction between the nSH2 and pTyr607 is highly unlikely to occur due to steric clashes, suggesting that the nSH2-pTYr607 interaction results in dimer formation. Figure 12 shows the most likely arrangement of a p85α-p85α homodimer (upper panel) and a p85α-p50α heterodimer (middle panel). Details of the nSH2-pTyr607 interaction in the context of the dimer are shown in Figure 12 (lower panel).

**Figure 12:**
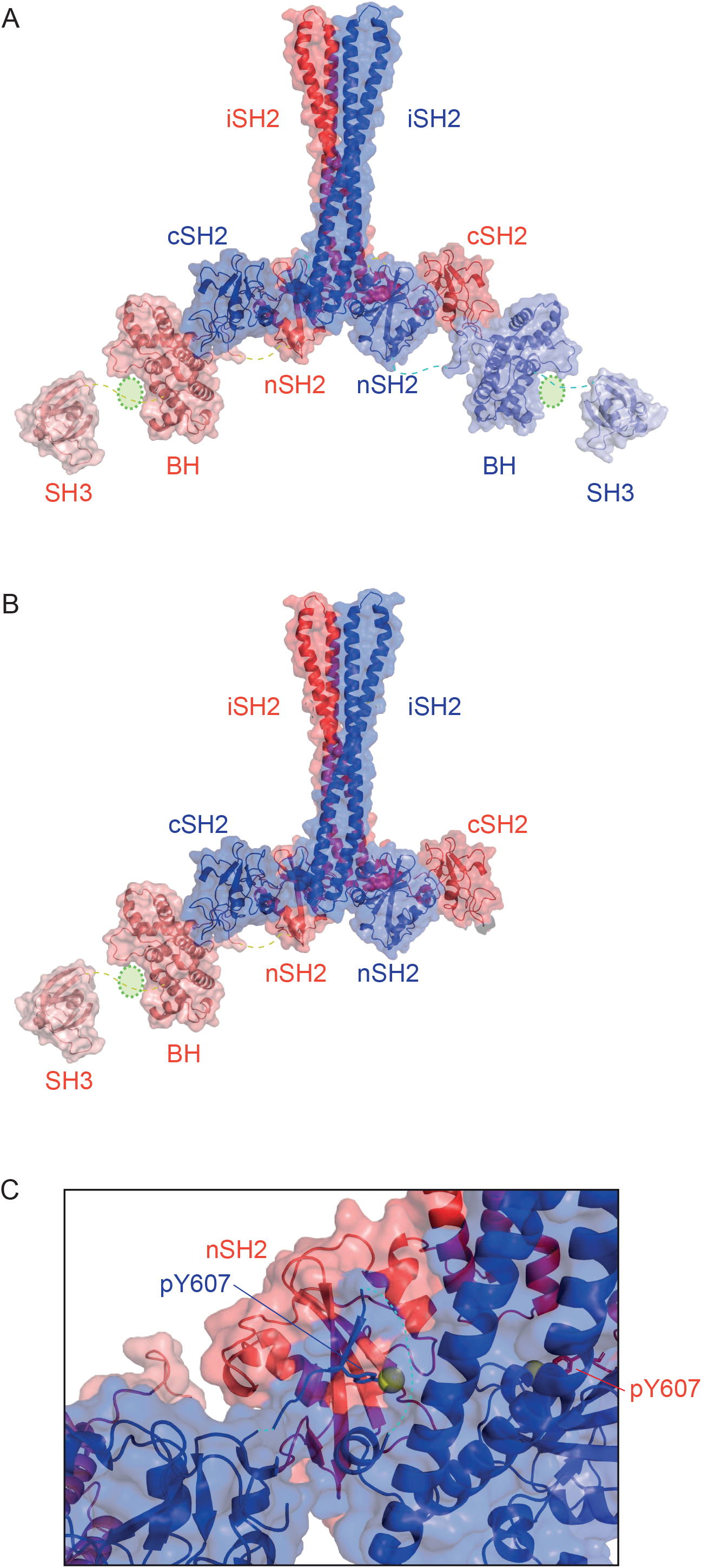
Structural model of the novel nSH2-pTyr607 mediated dimers. The model was constructed in Pymol using the following PDB codes: SH3 domain, 3i5r; BCR domain, 1pbw; nSH2-iSH2, 4l23; cSH2 5aul. The SH3, BH and nSH2 domains were rotated to minimize the distance between C- and N-terminal residues in adjacent domains. Domains were moved into one plane for clarity. The cSH2 was then positioned as follows: PDB 2iuh (p85 SH2 in complex with a c-Kit peptide) was aligned onto the model and the peptide component was mutated to the p85 sequence. The linker between the end of the iSH2 structure and the start of the peptide is 6 residues but there is no linker at all between the end of the peptide and the SH3 domain. When the iSH2 and SH3 domains were moved to be correctly positioned for the peptide to interact with the nSH2 it was not possible for an intramolecular nSH2-peptide interaction without steric clashes. A second monomer was constructed and the nSH2-peptide interaction used to derive the approximate positions of the domains in a dimer. (A) The potential p85α homodimer. The two chains of the dimer are shown in blue and red, with pale shades at the N-terminus and darker shades at the C-terminus. The domains are linked by dotted lines and the G protein binding site of the BH domain is indicated by a green circle. (B) The potential p85α -p50α heterodimer, colours as above except that p50 is red and p85 is blue. (C) Close up view showing the details of the intermolecular interaction between pTyr607 (blue) and the nSH2 (red). The sidechain of Tyr607 is shown as blue sticks and the phosphate group as a yellow sphere.

The p85 subunits are well documented to homodimerize via a *trans* interaction between the N-terminal SH3 domain and its juxtaposed proline rich motif (PR1) (41). The RhoGAP (BH) domain has also been shown to mediate homodimerization (41). Both of these interactions are relatively weak and it has been proposed that multiple sites of interaction may contribute to producing a stable p85 homodimer, allowing multiple levels of co-ordinated regulation that are common in signalling systems (40, 41). The pTyr607-nSH2 interaction we have identified provides a novel mechanism to mediate dimerization of the PI3K regulatory subunits via the C-terminal regions of the proteins (Figure 12). This interaction is high affinity and therefore sufficient to support dimerization in the absence of other contributing interactions. Unlike the SH3-PR1 and BH-mediated dimerization modes, both of which involve N-terminal regions of the protein, the pTyr607-nSH2 interaction also exists in the shorter isoform p50α and likely in p55α (Figure 12). Our data demonstrate that heterodimerization is supported between p85β and both p50α and p85α, but could potentially occur between any of the four phosphorylated regulatory subunit isoforms, as the nSH2 domains from p85α and p85β bind to phosphopeptides representing sequences from both p85α and p85β. The nSH2 and peptide sequences for p85α are identical to its splice variants p55α and p50α, implying that any combination of isoforms could potentially form heterodimers. Our affinity data indicate preference for binding to the p85β peptide in both cases, which is not surprising as the p85β peptide conforms well to the consensus sequence for a p85 SH2 domain binding target. This suggests that α isoforms should preferentially heterodimerize with p85β, while p85β would preferentially form homodimers. The functional ramifications of this remain to be investigated. Importantly, this new mechanism of dimerization is also regulated by phosphorylation, only being present in conditions where ACK is active. This might explain why it has not been identified in previous studies, especially those using recombinant proteins.

Dimerization of the regulatory subunits also has implications for the binding of other potential cellular ligands. Different p85 dimers would retain the ability to bind to a unique subset of cellular partners defined by the regions not involved in dimerization. p85 dimers mediated by N-terminal interactions could retain the ability to interact with, for example, the p110 catalytic subunit. Dimerization mediated by N-terminal regions also seems to be a pre-requisite for p85 to bind to PTEN and enhance its phosphatase activity, so negatively regulating PIP_3_ production in cells (40). Importantly a p85 dimer where the nSH2 domain is bound to pTyr607 *in trans* is unlikely to retain the ability to bind to the p110 catalytic subunits: multiple contacts to the p110 subunit occur within the nSH2 and iSH2 regions of the regulatory subunits. Analysis of the available structural data suggests that, if the nSH2 is bound to a phosphopeptide located in the iSH2, clashes would exist with the p110 catalytic subunit, implying that an pTyr607-nSH2 interaction would alter the p85-p110 heterodimer (Figure 12). As p85-free p110 is unstable and degraded quickly, this could contribute to the decrease in PIP_3_ levels we observe.

A p85 homo or heterodimer utilizing the nSH2 in a classical phospho-tyrosine interaction will also be modulated in its ability to bind RTKs and therefore potentially in competing with the PI3-Kinase heterodimer for activation. On the other hand, the cSH2 would still be available to interact with RTKs (Figure 12). p85 interactions with RTKs are complex, with the nSH2 and cSH2 having different affinities for different pTyr motifs (31, 42, 43). The availability of only the cSH2 might be expected to have variable effects on the ability of a pTyr607-nSH2 p85 dimer to bind to RTKs. However the cSH2 domain is often seen to have higher affinity for receptor sequences than the nSH2 (31, 42, 43). Thus competition could be the mechanism underpinning the decrease in PIP_3_ we observe. This newly identified pTyr607-nSH2 dimer could also retain the ability to bind to and activate PTEN, likewise leading to decreased levels of PIP_3_. Further work will be required to determine whether pTyr607 p85 retains the ability to form dimers that can interact with RTKs and/or PTEN.

The dimers we describe here would, in contrast to those previously reported, leave the N-terminal regions of p85α/β available for further interactions (Figure 12). Interestingly the only potential catalytic domain of p85α/β, the RhoGAP domain (BH domain) resides in the N-terminal regions of the proteins and the catalytic cleft on this domain is available to engage substrates (Figure 12). The proline-rich regions and RhoGAP/BH domain of p85 have been shown to regulate the actin cytoskeleton, with their deletion inhibiting lamellipodia formation (44). Cdc42-WASP controlled formation of filopodia is also regulated by p85 (45). Although the p85 RhoGAP domain binds to both Rac1 and Cdc42, GAP activity towards either small G protein, if it exists at all, is very low (46). Indeed, p85 appears to increase signalling from these small GTPases, in opposition to the traditional role of a GAP domain. The RhoGAP/BH domain is also thought to be a GAP for Rab proteins, including Rab5, albeit with equally low activity, and to therefore play a role in receptor endocytosis (46, 47). The SH3 domain of the p85 subunits interacts with many target proteins, which, in different contexts, functionally confer mechanisms to activate PI3-Kinase independent of the SH2 domains, inhibit PI3-kinase, regulate endocytosis or regulate the p85 protein itself (48). All of these interactions would be available in a pTyr607-nSH2 driven dimer and could be functionally altered in this new dimeric configuration (Figure 12).

A growing roster of PI3-Kinase independent functions for the p85 regulatory subunits exists. p85 plays roles in various stress response pathways, regulating p53, XBP-1 and BRD7 (33, 34, 36, 49, 50). Intriguingly the p85 regulatory subunits are also known to act independently of PI3-Kinase to potentiate c-Jun N-terminal kinase (JNK)-mediated insulin resistance. In this instance p85 acts as a negative regulator of insulin/PI3-Kinase signalling by increasing JNK activation. The activation of JNK in response to insulin is mediated by a Cdc42/MKK4 pathway and p85 seems to act by increasing levels of active Cdc42 (51). Our data add another layer of regulation to this network of feedback loops. As well as activating MKK4, Cdc42 will activate ACK, which in turn will increase p85 levels, potentially reinforcing JNK-mediated insulin resistance. This suggests a potential role for ACK in type 2 diabetes. It remains to be determined which other interactions and cellular processes a pTyr607-nSH2 p85 dimer can support and therefore what the overall impact of Cdc42/ACK signalling on p85 function will be.

ACK has been shown previously to shuttle between the cytoplasm and the nucleus in a Cdc42 dependent manner (35). Once in the nucleus it has a number of important roles in regulating transcription e.g. by influencing the androgen receptor and FoxO transcription factors (13, 52). However unlike BRD7, which binds to the iSH2 region and translocates p85 to the nucleus (36), our data support a model where ACK and p85 access the nucleus independently and only interact once in this compartment. Subsequently the dimers that form also appear to exist predominantly in the nucleus, suggesting a novel nuclear function for the regulatory subunits based on the new dimer configuration described here. Many of the roles of p85 in the nucleus revolve around regulation of transcription of genes that control cell proliferation or senescence, alongside protection of the cell from different forms of stress. We therefore suggest that ACK promotes cell proliferation, at least in part, through phosphorylation of free p85 in the nucleus. This results in a parallel increase in nuclear levels of p85 and subsequent formation of the pTyr607-nSH2 dimer, which may play a role in regulating gene transcription.

The work presented here, along with a growing body of literature describing the cellular effects of ACK, indicates that ACK regulates PI3-Kinase signalling at multiple nodes, suggesting that signalling through PI3-kinase is extremely important to the physiological functions of ACK. Our initial discovery that the p85 regulatory subunits were new targets for ACK led us to assume that ACK would act to relieve the inhibitory effects of the regulatory subunits, leading to activation of PI3-Kinase, based on the pro-proliferative activity of ACK. The reduction in PIP_3_ levels we observe are hard to reconcile with the growth promoting activity of ACK. Our initial hypothesis was however always somewhat perplexing, as ACK can interact with and phosphorylate Akt directly, providing a potential membrane localization signal to contribute to the activation of Akt (18). Phosphorylation of Tyr176 also appears to drive Akt to the nucleus where it can phosphorylate the FoxO transcription factors, leading to their relocation to the cytoplasm and stimulation of the cell cycle. The reason why two pathways to Akt activation, *i.e.* RTKs/PI3-Kinase or ACK, are employed in cells seemed elusive(18). However our discovery that the ACK expression results in a decrease in PIP_3_ levels may go some way to explaining the necessity for direct Akt activation by ACK, independent of PI3-Kinase.

Our data implies that the effects that ACK is able to exert via the free p85 pool must be paramount to its pro-proliferative role, such that it also has the ability to maintain downstream activation of the canonical ‘PI3-Kinase’ pro-proliferative pathways (via Akt), while directing p85 to perform other vital roles in the nucleus. Via this mechanism, ACK would be able to shape PI3-Kinase signalling responses, promoting the activity of pathways downstream of Akt, whilst suppressing signalling in other PI3-Kinase effector pathways and stimulating free dimeric p85 functions (Figure 13). It remains to be investigated whether ACK can directly activate further pathways conventionally downstream of PI3-Kinase/PIP_3_.

**Figure 13:**
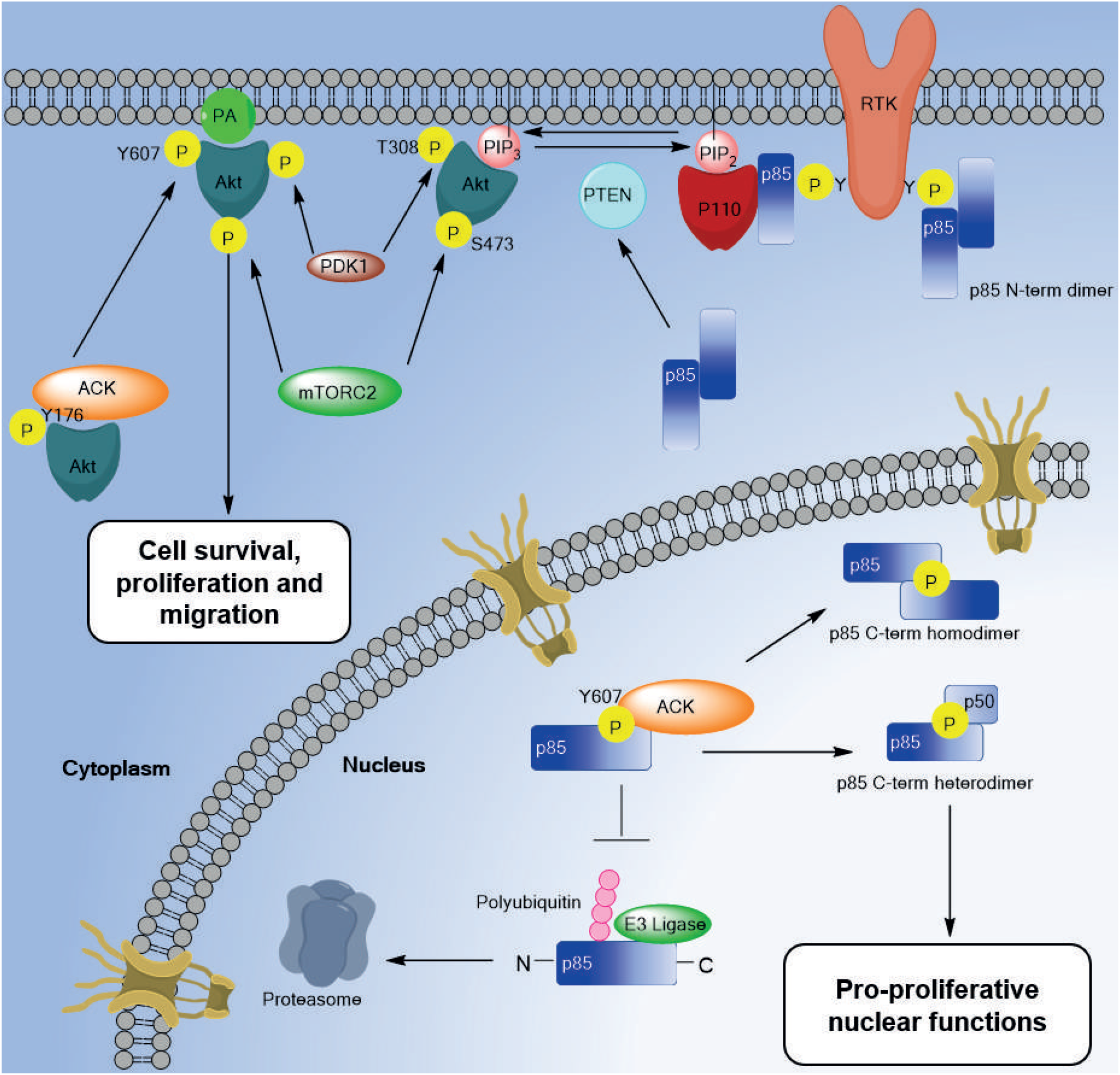
Model depicting the multiple roles of ACK in the regulation of PI3-Kinase signalling. ACK interacts with the PI3-Kinase regulatory subunit(s) (p85) in the nucleus, phosphorylating Tyr607. This leads to stabilization of p85 levels by preventing ubiquitination and subsequent degradation by the proteasome. Increased levels of p85 and pY607 promote dimerization of the regulatory subunits. We propose that this new dimeric conFigureuration of the regulatory subunits possesses pro-proliferative nuclear functions. The interaction between ACK and the regulatory subunits ultimately leads to a suppression PIP_3_ levels in the cell, however, ACK is able to promote cell proliferative signals by phosphorylating Akt, contributing to its activation. PDK1: Phosphoinositide-dependent kinase-1, mTORC2: mTOR complex 2, PTEN: phosphatase and tensin homologue, PA: phosphatidic acid.

Growing evidence in the literature demonstrates that the relative amounts of p85/p110 heterodimer and p110-free p85 homodimer are critical in regulating the PI3-Kinase pathway and a number of other pathways regulated by p85 homodimers. We demonstrate here that ACK perturbs this gatekeeper equilibrium and suggest that this feature of ACK contributes to the oncogenic potential of this kinase. Most importantly we have identified a new form of PI3-Kinase regulatory subunit dimers, which form exclusively in the nucleus and must undertake pro-proliferative roles.

## Abbreviations

BH: Bcr homology
CRIB: Cdc42/Rac interactive binding
EBD: EGFR binding domain
EGF: epidermal growth factor
GAP: GTPase activating protein
NES: Nuclear export signal
PIP_2_: phosphatidylinositol 4,5-bisphosphate
PIP_3_: phosphatidylinositol (3,4,5)-trisphosphate
RTK: receptor tyrosine kinase
SAM: sterile alpha motif
SH2: Src homology 2 domain
SH3: Src homology 3 domain
UBA: ubiquitin association

## Author contributions

H.R.M. and D.O. conceptualized the project. N.S.C., M.F., J. J. V-G., C.M.S., T.D.L., J.I.W., J.C., K.K., Q.Z., M.J.O.W., M.J.B.B., C.C., H.R.M and D.O. designed and performed the experiments. N.S.C., M.F., C.M.S., J.I.W., J.C., Q.Z., M.J.O.W., M.J.B.B., C.C., H.R.M and D.O. interpreted and analysed the data. N.S.C., M.F., H.R.M and D.O. wrote the manuscript with contributions from all other authors.

## Funding

This research was supported by: BBSRC DTG/A studentships (BB/F017464/1 and BB/A517685/1) to NSC and JV-G; an MRC iCASE (MR/N018354/1) to DO and CC; a Churchill Scholarship (awarded by the Winston Churchill Foundation of the United States) to CS; a CR-UK (C4750/A19013) programme grant to TDL; a National Overseas Scholarship, Government of India and a Cambridge Commonwealth Scholarship to KK; a BBSRC research grant (BB/P013384/1) to MJOW and a CR-UK project grant (C11309/A5148) to DO and HRM. Thanks also to the Amgen Foundation for the financial support provided through the Amgen Scholars Programme to JC.

### Acknowledgements

We are very grateful to Dr Marc de la Roche and Dr Catherine Lindon for helpful discussions. We are also indebted to Dr Elizabeth Smethurst for expert assistance in quantifying PIP_3_ levels. Phosphorylation site mapping was carried out by Dr. Mike Deery, Cambridge Centre for Proteomics, University of Cambridge.

## Supplementary Material

**Figure S1:**
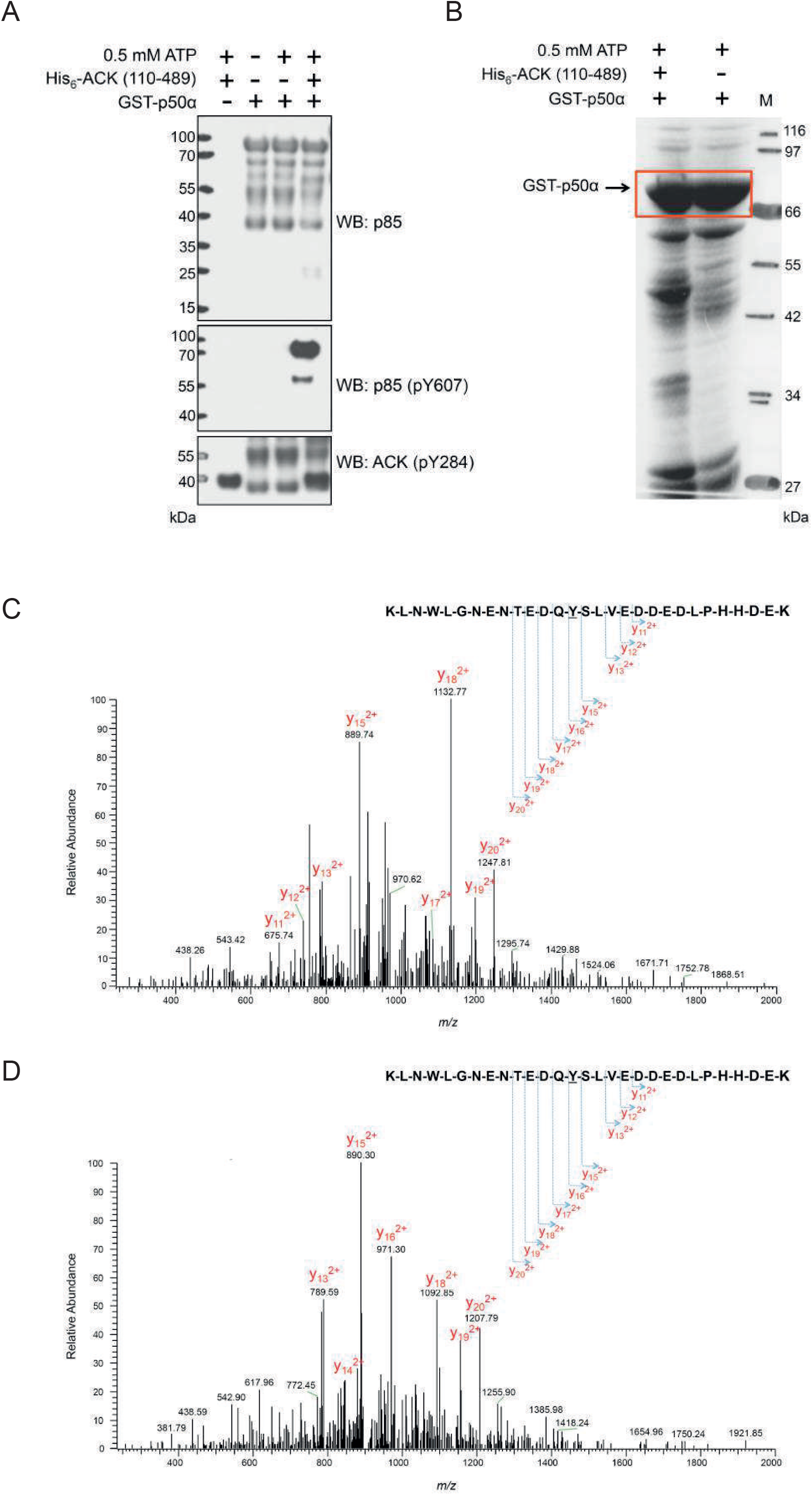
Identification of ACK phosphorylation sites in GST-p50α. (A) GST-p50α is phosphorylated on Y307 by His6-ACK (110-489) *in vitro*. Full-length GST-p50α was purified from *E. coli* and incubated at 30 °C for 30 minutes with His6-ACK (110-489) purified from Sf9 cells. Reactions were analysed by SDS-PAGE and the phosphorylation of GST-p50α at Y307 was assessed by western blotting using an antibody raised against the sequence (D-Q-Y(p)-S-L) in human p85α. The presence of autophosphorylated His6-ACK (110-489) was confirmed by western blotting using an anti-ACK (pY284) antibody. (B) *In vitro* kinase assay reactions for GST-p50α in the presence and absence of His6-ACK (110-489) were resolved on a NuPAGE 4-12% gel and proteins were visualized by Coomassie staining. (C) Phosphorylation sites in excised full-length GST-p50α (red box, panel B) were identified by LC-MS/MS. Annotated MS/MS spectrum of the tyrosine phosphorylated peptide KLNEWLGNENTEDQYSLVEDDEDLPHHDEK (m/z 923.65, 4+). The doubly charged C-terminal y fragment ions are highlighted which specifically represent the tyrosine phosphorylation. This can be compared with the non-phosphorylated peptide (D) (m/z 903.65, 4+) in which the m/z values are 40 units lower (corresponding to a mass of 80 Da) from y16, which is indicative of tyrosine phosphorylation.

**Figure S2:**
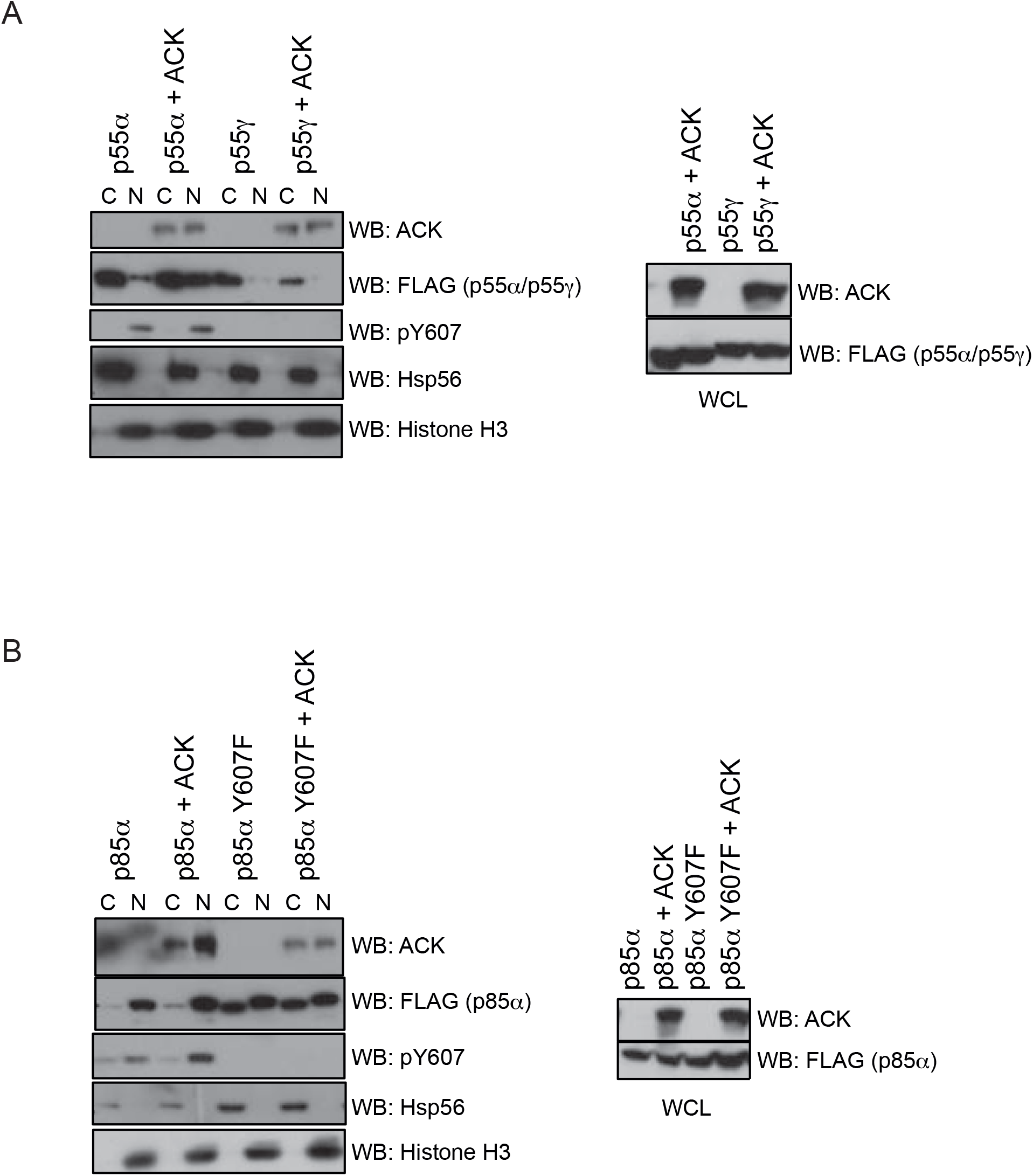
(A) wt p50α and wt p55γ were expressed alone or co-expressed with ACK in HEK293T cells. Cells were lysed and separated into cytoplasmic and nuclear-enriched fractions. Fractions were analysed by western blotting using the antibodies indicated (left hand panels). Fractionation was judged using anti-Hsp56 to identify cytoplasmic extracts and anti-Histone H3 to identify nuclear-enriched fractions. Samples of whole cell lysates (WCL) were analysed by western blotting, to show total levels of proteins, using the antibodies indicated and are shown in the right hand panels. Results are representative of at least three independent experiments. (B) wt p85α and Y607F p85α were expressed alone or co-expressed with ACK in HEK293T cells. Cells were lysed and separated into cytoplasmic and nuclear-enriched fractions. Fractions were analysed by western blotting using the antibodies indicated (left hand panels). Fractionation was judged using anti-Hsp56 to identify cytoplasmic extracts and anti-Histone H3 to identify nuclear-enriched fractions. Samples of whole cell lysates (WCL) were analysed by western blotting, to show total levels of proteins, using the antibodies indicated and are shown in the right hand panels. Results are representative of at least three independent experiments.

